# Genomic assessment of local adaptation in dwarf birch to inform assisted gene flow

**DOI:** 10.1101/727156

**Authors:** James S. Borrell, Jasmin Zohren, Richard A. Nichols, Richard J. A. Buggs

## Abstract

When populations of a rare species are small, isolated and declining under climate change, some populations may become locally maladapted. Detecting this maladaptation may allow effective rapid conservation interventions, even if based on incomplete knowledge. Population maladaptation may be estimated by finding genome-environment associations (GEA) between allele frequencies and environmental variables across a local species range, and identifying populations whose allele frequencies do not fit with these trends. We can then design assisted gene flow strategies for maladapted populations, to adjust their allele frequencies, entailing lower levels of intervention than with undirected conservation action. Here, we investigate this strategy in Scottish populations of the montane plant dwarf birch (*Betula nana*). In genome-wide single nucleotide polymorphism (SNP) data we found 267 significant associations between SNP loci and environmental variables. We ranked populations by maladaptation estimated using allele frequency deviation from the general trends at these loci; this gave a different prioritization for conservation action than the Shapely Index, which seeks to preserve rare neutral variation. Populations estimated to be maladapted in their allele frequencies at loci associated with annual mean temperature were found to have reduced catkin production. Using an environmental niche modelling (ENM) approach, we found annual mean temperature (35%), and mean diurnal range (15%), to be important predictors of the dwarf birch distribution. Intriguingly, there was a significant correlation between the number of loci associated with each environmental variable in the GEA, and the importance of that variable in the ENM. Together, these results suggest that the same environmental variables determine both adaptive genetic variation and species range in Scottish dwarf birch. We suggest an assisted gene flow strategy that aims to maximize the local adaptation of dwarf birch populations under climate change by matching allele frequencies to current and future environments.

## Introduction

Climate change is predicted to become a major driver of global biodiversity loss (Bellard et al., 2012; Urban, 2015). Species that lack relevant phenotypic plasticity (Gratani, 2014; Nicotra et al., 2010) may survive environmental changes by dispersing to new locations, consequently tracking conditions they are currently adapted to (Aitken et al. 2008; Meier et al. 2012), or remaining in the same location and rapidly evolving adaptation to their new environments from standing genetic variation or gene flow (Aitken et al., 2008; Alberto et al., 2013). Migration in response to rapid climate change may be particularly difficult for plants (Corlett and Westcott, 2013; Hampe and Petit, 2005; Zhu et al., 2012). In some cases plants lack the dispersal ability to keep pace with accelerated climate shifts (Loarie et al., 2009), there is an absence of potential habitat at higher latitudes (McKenney et al., 2007) and altitudes (Engler et al., 2011), or suitable new habitats may be separated by too large distances (Meier et al., 2012). In these cases, conservation managers aiming to prevent extinction of species or populations face a choice between relying on *in situ* evolution to track the environmental change, or attempting conservation interventions such as assisted migration or assisted gene flow that seeks to enable, facilitate or accelerate adaptation.

To evaluate whether interventions are appropriate, a first step is understanding current local adaptation and the potential for adaptation to future environments (Davis et al., 2005; Hoffmann et al., 2017; Funk et al 2019). The classical way to identify local adaptation is via reciprocal transplant experiments (Kawecki and Ebert, 2004; Leimu and Fischer, 2008; Pardo-Diaz et al., 2015). However, this approach is often unfeasible for wild organisms with long generation times in need of urgent conservation, meaning that more rapid approaches using genomics are desirable (Williams et al., 2008).

Genotype-environment association (GEA; also referred to as environmental association analysis, EAA) methods are increasingly used to identify loci involved in local adaptation (Abebe et al., 2015; Ahrens et al., 2018; Bay et al., 2017; Coop et al., 2010; Flanagan et al., 2018; Günther and Coop, 2013; Rellstab et al., 2015; Funk et al 2019). These approaches detect replicated signatures of selection (SNPs that deviate strongly from estimated neutral population structure) across many independent populations. Thus far the majority of studies to apply GEA in tree species have been targeted at candidate genes, and surveyed fewer than 350 loci (Keller et al., 2012; Nadeau et al., 2016; Rellstab et al., 2016; Wang et al., 2016).

Building on the assumption that GEA captures an important component of locally adaptive allelic variation, especially if based on genome-wide markers, we may extend it to rapidly assess local adaptation and adaptive potential within populations. The principal of this approach is the detection of discordance between genotype and environment, in certain populations, as an indicator of reduced local adaptation and vulnerability to future demographic decline (Alberto et al., 2013). In a previous study, Rellstab *et al*. (2016) developed a model to estimate the average change in allele frequency at environmentally-associated loci that would be required to respond to projected future environmental conditions. They based this estimate on the allele frequency changes that would maintain the present-day associations between genotype and environment and term this mismatch, the risk of non-adaptedness (RONA). For clarity we term this ‘future risk of non-adaptedness’ (f-RONA) and comment that rather than a ‘risk’ this is a forecast, but for consistency we maintain the same terminology in this manuscript. This approach to estimating adaptation has many simplifying assumptions. Environmental variation in nature is complex, as are the mechanisms by which organisms adapt to them, but as Funk et al (2019) argue, any available evidence may improve conservation decision making.

Here, we extend the work of Rellstab *et al*. (2016) to explicitly define c-RONA, the ‘current risk of non-adaptedness’, that is the average change in allele frequency at climate-associated loci required to match our estimate of the optimum for current climatic conditions (for a given environmental factor). Current risks are likely to be particularly important for species that are already declining due to climate change, and have small isolated populations. Furthermore, we extend the univariate RONA model to a multi-locus analysis of genome-wide markers, and use best linear unbiased prediction (BLUP) to improve our estimate of the effect of each allele.

In populations where c-RONA is high, local genotypes would not match local environmental variables as expected. Therefore, a possible management intervention is to use assisted gene flow (AGF) to introduce more appropriate alleles or adjust population allele frequencies. Here, AGF is defined as the managed movement of individuals or gametes between populations, from source populations that have been selected with the aim of accelerating adaptation, so that it is faster than would occur by passive natural dispersal alone (Aitken and Whitlock, 2013). This AGF strategy could be used to inform sourcing of seed stock for reforestation programs (Boshier et al., 2015) and mitigate maladaptation to future climate (Aitken and Bemmels, 2016; Havens et al., 2015; Jin et al., 2016). Importantly, only modest translocation of genotypes may enhance adaptation by introducing genetic variation upon which selection can act to further refine local allele frequencies (Bay et al., 2017; Pavlova et al., 2017). Conversely such interventions could have negative effects (i.e. outbreeding depression) if they cause gene flow between populations with undetected adaptive differentiation (Frankham et al., 2011; Pavlova et al., 2017). We note that where target populations are small, maladapted and dominated by drift, Assisted Gene Flow is equivalent to Genetic Rescue (see Aitken and Whitlock (2013) for a detailed review).

If AGF is to be effective, there must be appropriate populations from which to source migrants. Such populations might be found towards the species’ retreating range edge or other locations where environmental conditions are closer to those anticipated in the future (Olson et al., 2013). To design a sampling strategy that encompasses both environmental gradients and declining range edge populations threatened by environmental change, we can use environmental niche models (ENMs) (Maguire et al., 2015). ENMs project the distribution of species’ ranges under current and future climate scenarios based on observation data and can guide effective sampling (Elith and Leathwick, 2009). ENMs are also an established tool for conservation practitioners seeking to understand major climatic selection pressures and projected range shifts for threatened species, but often lack integration and comparison with genomic assays of local adaptation (Hällfors et al., 2015; Razgour et al., 2019).

Here, we conduct GEA and ENM analysis of wild populations of dwarf birch (*Betula nana*), for which we have field observation and genome-wide population genetic data. In the UK, dwarf birch is a nationally scarce montane tree that has experienced an accelerated decline in recent decades, likely due to the combined impact of anthropogenic climate change and moorland management that permits over-browsing and burning (Aston, 1984; Borrell et al., 2018; Wang et al., 2014; Zohren et al., 2016). Dwarf birch, like many tree species, is the focus of a conservation program to restore populations, delimit management units and prioritise the protection of important genetic diversity (Koskela et al., 2013). Germplasm collection from central Scottish Highland populations is already underway for reintroduction to other parts of the species former range (pers. obs. J Borrell). Previous research by our group has found that despite extensive fragmentation, most populations of dwarf birch in the UK contain diversity comparable to that of large, unfragmented Scandinavian populations (Borrell et al., 2018). Nevertheless, we concluded that this diversity has become increasingly partitioned among populations. In other words, much of the adaptive diversity in dwarf birch is still extant in the UK, but due to restricted gene flow and dispersal, marginal populations may be maladapted due to a failure to track environmental change, or by drift of adaptive alleles away from their optimum frequency. There is limited potential for naturally occurring gene flow to enhance future adaptation in many populations.

In species subject to conservation management such as dwarf birch, evolutionary processes have sometimes been overlooked, despite the importance of adaptation to species persistence (Eizaguirre and Baltazar-Soares, 2014; Fitzpatrick and Keller, 2015). Therefore the adaptive potential of populations may be underrepresented in conservation prioritization strategies (Funk et al., 2019; Harrisson et al., 2014). For example, where genetic diversity information is available to conservationists, metrics that score populations on neutral genetic distinctiveness, such as the Shapley Index are often used (Haake et al., 2007; Isaac et al., 2007; Volkmann et al., 2014). However there is no guarantee that neutral and adaptive diversity will be correlated (Bonin et al., 2007), and indeed approaches designed solely to promote or conserve neutral diversity may be harmful (Reed and Frankham, 2003; Weeks et al., 2016). Therefore evaluating adaptive diversity, rather than using more established metrics of genetic diversity should improve the prioritisation decisions in species management, though see Kardos and Shafer, (2018) for potential pitfalls.

To explore potential management strategies for dwarf birch, that takes into account local adaptation and evolutionary potential, we first characterise the species’ range using ENMs under present and projected future climate scenarios. We evaluate these ENMs by assessing whether populations on the margins of the inferred distribution had lower scores for phenotypic and fitness proxies for local adaptation. Second, we use GEA to survey putative adaptive loci across the species’ range and estimate c-RONA to identify populations with a discordance between genotype and environment. The combined ENM and GEA data present an opportunity to test the hypothesis that limiting environmental variables (which have higher discriminatory power in an ENM) have more genomic loci associated with them in GEA, perhaps as a result of stronger selection for adaptation (an alternative would be that certain variables limit species’ ranges precisely because they lack genetic adaptation). We provide preliminary evidence in support of this hypothesis in dwarf birch. Third, we evaluate our estimates of non-adaptedness (c-RONA) of dwarf birch populations against the Shapley Index, an existing conservation prioritization most often applied to neutral markers. Finally, we illustrate a strategy of AGF to maximize adaptive genetic diversity and hence sustain the adaptive potential of British dwarf birch populations. We discuss the advantages and limitations of this approach in the context of managing dwarf birch and other plants exposed to rapid environmental change.

## Methods

### Environmental niche modelling

To determine the environmental variables influencing the present and future distribution of dwarf birch in the UK, we developed an ENM based on 763 resampled fine-scale (≤1 km) records from the period 1960-present. Records were sourced from national databases, conservation partners and fieldwork observations (see Borrell *et al*. 2018). Nineteen bioclimatic layers were obtained from the WorldClim database (www.worldclim.org) at 1km resolution (Hijmans et al., 2005), for the period 1960-1990, including 11 temperature and eight precipitation derived variables reflecting annual trends, seasonality and limiting environmental factors. High resolution elevation data was used to compute slope and aspect terrain characteristics using the *Raster* package (Hijmans & Etten, 2012) in R software (R Development Core Team, 2014). These variables are indicators of soil moisture, erosion, wind and solar radiation (Hoersch et al., 2002). To avoid overfitting, we removed multiple highly correlated variables (correlation coefficient >0.7), retaining 10 for analysis (preferring less derived, e.g. Annual Mean Temperature, rather than Monthly or Quarterly values) (Table 1, Figure S1). Elevation was excluded due to its high correlation with temperature (Parolo et al., 2008). Temperature was retained because it captures the projected change in climate change models, whilst elevation does not. All retained variables were standardized to a mean of zero and unit variance. Eight further datasets consisting of the same retained variables were generated under four representative concentration pathways (RCP) defined by the Intergovernmental Panel on Climate Change Fifth Assessment (IPCC, 2014a) at each of two future time points (2045-65 and 2081-2100). These projections allow estimation of future temperature and precipitation values across the study area derived from the Community Climate System Model (Gent et al., 2011) (Table S1).

**Table 1.** Contribution of retained environmental variables to the dwarf birch environmental niche model (ENM), and the number of environmentally associated loci detected.

| Variable | Description | Correlated Variables<sup>1</sup> | ENM percent GEA Loci contribution<sup>2</sup> |  | GEA Loci (inc. cor.)<sup>3</sup> |
| --- | --- | --- | --- | --- | --- |
| AMTemp | Annual Mean Temperature | MTColdQ, MTColdM | 34.9 | 17 | 64 |
| MTWarmM | Max Temperature of Warmest Month | MTWarmQ | 22.1 | 2 | 6 |
| MDR | Mean Diurnal Range | - | 14.8 | 71 | 71 |
| ISO | Isothermality | - | 14.6 | 11 | 11 |
| APrec | Annual Precipitation | PColdQ, PWetM, PSeason, PWetQ, PWarmQ, PDryM, PDryQ | 7.3 | 2 | 21 |
| Slope | Slope | - | 2.8 | 7 | 7 |
| MTDryQ | Mean Temperature of Driest Quarter | - | 1.6 | 7 | 7 |
| TS | Temperature Seasonality | ATempR | 1.4 | 1 | 3 |
| MTWetQ | Mean Temperature of Wettest Quarter | - | 0.3 | 7 | 7 |
| Aspect | Aspect | - | 0.2 | 4 | 4 |

<sup>1</sup> Correlated variables include Mean Temperature of the Coldest Quarter (MTColdQ); Minimum Temperature of the Coldest Month (MTColdQ); Mean Temperature of Warmest Quarter (MTWarmQ); Precipitation of Coldest Quarter (PColdQ); Precipitation of Wettest Month (PWetM); Precipitation Seasonality (Pseason); Precipitation of Wettest Quarter (PWetQ); Precipitation of the Warmest Quarter (PWarmQ); Precipitation of Driest Month (PDryM); Precipitation of Driest Quarter (PDryQ); Annual Temperature Range (ATempR).
<sup>2</sup> Percentage contribution is calculated as the increase in regularized gain added to the contribution of the corresponding variable over each iteration of the model.
<sup>3</sup> Total number of SNPs associated with both the retained variable, as well as related highly correlated variables that were excluded from the ENM analysis.

The ENMs were generated using MaxEnt (Phillips et al., 2006) within the *dismo* package (Hijmans et al., 2011). We performed 50 randomly subsampled replicate runs with 25% of observations retained for cross-validation. Models were further evaluated using a binomial test of omission rate and Area Under the Receiver Operating Characteristic Curve (AUC). A species occurrence threshold to assess changes in occupied area was defined by ‘maximum training sensitivity plus specificity’, which optimizes the trade-off between commission and omission errors (Liu et al., 2016). Rank and percentage contribution of environmental variables is reported here, as these have been demonstrated to capture biologically important factors (Searcy & Shaffer, 2016).

### Phenotypic data and habitat suitability projections

We identified 29 dwarf birch populations that encompass the extant UK range (Table 2, Figure S2). To test the performance of our ENM, we collected extensive phenotypic measurements of traits related to reproductive output and fitness in 20-30 individuals per population in June-August 2013. These included: the number of male and female catkins, plant area, plant height and diameter of the largest stem. Cambial tissue samples were retained for genetic analysis. A subset of 18 populations was also tested for seed viability in germination experiments, a fitness proxy relevant to population persistence (Alsos et al., 2003). Seed were collected in late summer, over-wintered at 4°C then kept in moist conditions at 18-20°C with a 14h photoperiod for 60 days the following spring. For nine of these populations, 100-day survival of seedlings during the following Spring was measured (See Supplementary Materials for details).

**Table 2.** Summary information for 29 dwarf birch populations, including the number of genotyped and phenotyped individuals, habitat suitability (HSI).

| Location | Pop. | Lat. | Long. | Elev. (m) | Genotyped | Phenotyped | HSI | c-RONA | Shapley<sub>NEUTRAL</sub> |
| --- | --- | --- | --- | --- | --- | --- | --- | --- | --- |
| Ben Loyal | BL | 58.4 | -4.4 | 300 | 6 | 30 | 0.38 | 0.194 | 0.011 |
| Meall Odhar | MO | 58.16 | -4.42 | 404 | 6 | 29 | 0.45 | 0.168 | 0.006 |
| Beinn Enaiglair | BE | 57.79 | -5.01 | 480 | 5 | 27 | 0.37 | 0.479 | 0.01 |
| Luichart | LH | 57.72 | -4.9 | 268 | 6 | 29 | 0.54 | 0.131 | 0.008 |
| Ben Wyvis W | BW | 57.65 | -4.6 | 482 | 5 | 30 | 0.77 | 0.149 | 0.01 |
| Ben Wyvis E* | DG | 57.65 | -4.56 | 472 | - | 21 | 0.75 | - | - |
| Loch Meig | ME | 57.53 | -4.8 | 450 | 6 | 26 | 0.57 | 0.128 | 0.005 |
| Glen Cannich | GC | 57.34 | -4.86 | 455 | 6 | 31 | 0.51 | 0.045 | 0.027 |
| Faskanyle* | FS | 57.33 | -4.85 | 486 | - | 17 | 0.66 | - | - |
| Dundreggan Excl. | DE | 57.23 | -4.75 | 448 | 6 | 30 | 0.81 | 0.174 | 0.009 |
| An Suidhe | AS | 57.22 | -4.81 | 661 | 2 | 17 | 0.77 | 0.219 | 0.119 |
| Beinn Bhreac | BB | 57.21 | -4.82 | 500 | 6 | 33 | 0.66 | 0.366 | 0.008 |
| Portclair | PC | 57.2 | -4.64 | 478 | 6 | 38 | 0.54 | 0.081 | 0.008 |
| River Avon | AV | 57.14 | -3.49 | 549 | 6 | 28 | 0.59 | 0.306 | 0.01 |
| Monadhliaths | MD | 57.06 | -4.31 | 712 | 6 | 6 | 0.49 | 0.222 | 0.01 |
| Meall an tslugain | SL | 57.05 | -3.45 | 633 | 6 | 31 | 0.59 | 0.085 | 0.035 |
| Loch Muick E | MU1 | 56.92 | -3.2 | 492 | 6 | 31 | 0.17 | 0.223 | 0.006 |
| Loch Muick W | MU2 | 56.92 | -3.21 | 517 | 6 | 16 | 0.1 | 0.218 | 0.008 |
| Loch Laggan | LG | 56.89 | -4.54 | 364 | 6 | 33 | 0.35 | 0.064 | 0.007 |
| Loch Loch | LL | 56.85 | -3.65 | 673 | 6 | 32 | 0.57 | 0.106 | 0.005 |
| Ben Gullabin | BG | 56.84 | -3.47 | 594 | 1 | 7† | 0.58 | 0.194 | 0.422 |
| Loch Rannoch | LR | 56.76 | -4.42 | 499 | 6 | 28 | 0.23 | 0.097 | 0.008 |
| Rannoch West | RW | 56.65 | -4.79 | 306 | 6 | 32 | 0.61 | 0.218 | 0.007 |
| Rannoch Moor B | RB | 56.6 | -4.74 | 304 | 6 | 10 | 0.51 | 0.169 | 0.008 |
| Rannoch Moor A* | RA | 56.6 | -4.74 | 295 | - | 27 | 0.51 | - | - |
| Lennox | LX | 55.97 | -4.28 | 164 | 2 | 10 | 0 | 0.241 | 0.102 |
| Emblehope† | EM | 55.24 | -2.48 | 448 | 1 | 1† | 0.06 | 0.254 | 0.155 |
| Spadeadam† | SA | 55.05 | -2.57 | 275 | 1 | 1† | 0.01 | 0.321 | 0.35 |
| Teesdale† | TD | 54.65 | -2.28 | 499 | 2 | 2† | 0.06 | 0.291 | 0.133 |

\*Populations not submitted for genetic analysis, but are considered in the comparison of HSI and reproductive output.
†Populations were exhaustively sampled.

To assess change in habitat quality across the study area, we first plotted the ENM derived habitat suitability index (HSI) estimates for all populations under current and future conditions. Second, ENM performance was assessed using a generalized linear model with a quasipoisson error distribution to test for a relationship between present time HSI estimates and mean population catkin counts. We also tested for a relationship between HSI (explanatory variable) and mean germination rates (response variable) using a quasibinomial error distribution. Here we are explicitly testing the hypothesis that plants displayed greater reproductive output in locations with a higher ENM derived HSI.

### RAD sequencing

The genetic samples used in this study are a subset of those described in (Borrell et al., 2018). Briefly, DNA was extracted from 130 individuals (Table 2) and submitted to Floragenex (Oregon, USA) for 100bp single-end RAD sequencing with the enzyme *PstI*. Raw reads were filtered using Stacks v1.35 (Catchen et al., 2013) and aligned to the dwarf birch genome, retaining only reads that align uniquely (Wang et al., 2013) using Bowtie2 (Langmead and Salzberg, 2012) and the *ref_map.p*l pipeline. SNPs were called with a minimum depth of 5, the bounded model and a minimum log likelihood of −20, with corrections made using *rxstacks*. Finally, we filtered for loci present in ≥8 populations, and a minor allele frequency >0.05.

### Genomic signatures of local adaptation

We first used BayeScan (Foll and Gaggiotti, 2008) to compare allele frequency differences among populations and identify F_ST_ outlier loci. Analysis was performed with 50,000 iterations thinned every 10, with 20 pilot runs, a burn-in of 50,000 iterations and other parameters at default. Whilst F_ST_ outliers are candidate loci of adaptation, they can also emerge because of selection due to deleterious alleles, hybrid zones and historical demography (Bierne et al., 2013). Thus, we use relaxed BayeScan parameters to screen outlier loci prior to GEA analysis in Bayenv2 (Günther and Coop, 2013).

Bayenv2 incorporates neutral genetic structure using a covariance matrix based on neutral markers and attempts to identify correlations between outliers and environmental gradients, potentially reducing false positives (De Mita et al., 2013). Based on recommendations in François *et al*. (2016), to further minimize false positives we initially excluded loci detected in BayeScan to compute a null covariance matrix of relatedness between populations, over 100,000 iterations and five independent runs. We then tested all loci (including those initially identified by BayeScan) under an alternative model where allele frequencies are determined by a combination of the covariance matrix and an environmental variable. We performed our analysis independently across all environmental variables, with the expectation that correlated predictors would return subsets of the same markers. The posterior probability that a locus is under selection, across each independent environmental variable was assessed using Bayes factors (BF), with log10 posterior odds ratio values >1 defined as strong support (Jeffery, 1961). We averaged BFs over independent runs as recommended by Blair et al. (2014), and following Günther & Coop (2013) we retained loci as good candidates if, in addition to a high BF, they also fell in the top 10% of Spearman correlation coefficient values, to further reduce false positives. For comparison, we also independently tested for signatures of local adaptation using Redundancy Analysis (RDA) (Forester et al., 2018; Rellstab et al., 2015), (see Supplementary Materials) though we consider only the candidates identified using Bayenv2 in subsequent analyses.

### Gene expression

To provide an additional line of evidence on the activity of our candidate adaptive loci, we extracted up to 10,000bp flanking each side of the candidate locus from the *B. nana* reference genome and searched for these sequences in an RNA expression database using dwarf birch tissues derived from our genome reference plant under glasshouse conditions (Wang et al., 2013). Briefly, RNA was extracted from fresh dwarf birch leaves and flowers using a modified RNAeasy Plant Mini Kit (Qiagen, Hilden, Germany), incorporating additional CTAB and phenol-chloroform steps to generate 100bp paired-end reads with an average insert size of 280bp (for full methods see Zohren, 2016). These were mapped to the reference genome using Trinity software (Grabherr et al., 2013).

### Maladaptation under present and future conditions

We carried out RONA analysis on the nine standardized environmental variables that were associated with six or more candidate loci, allocating each locus to the single environmental variable with the largest Bayes factor (thereby avoiding double-counting a locus in the c/f-RONA calculations below). We estimated the vector of effect sizes, ***β***, in which each row corresponds to a locus, using R package rrBLUP (Endelman 2011). In this analysis, the vector of allele frequencies ***f*** for each population was used as the predictor of the environment in that location. The sum of ***fβ*** gives an estimate of the environment (the value of the environmental variable) to which the population would be best adapted. The residual deviation of the observed value from this expectation is a measure of the deviation from the optimum environment for that population (c-RONA), and is proportional to the change in allele frequency that would be required to match the population to its local environment (weighted by ***β***). This measure is therefore analogous to those employed by Rellstab *et al*. (2016) and Pina-Martins *et al*. (2018), which quantify the mismatch between genotypes and environment in terms of allele frequencies. We combined information across variables by calculating the mean of the absolute residuals. Similarly, we could calculate the difference from the projected values of the environmental values under each climate change scenario to estimate f-RONA (Figure 2).

**Figure 1.**
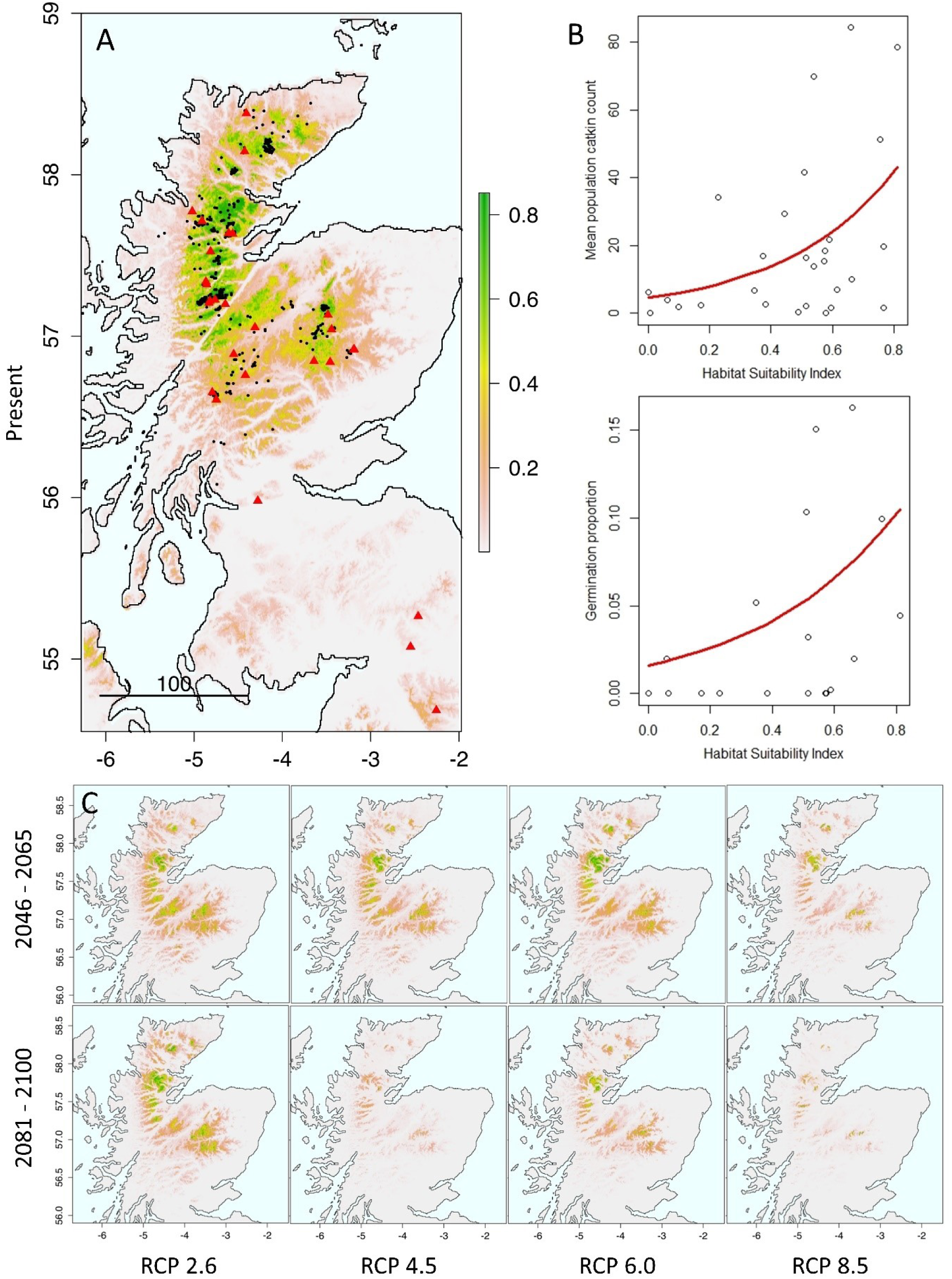
A) Environmental niche model of dwarf birch habitat suitability (HSI) under current environmental conditions, black points indicate species distribution records and red points indicate sampled locations included in this study. B) Regression of phenotypic fitness traits against the derived habitat suitability index. C) dwarf birch habitat suitability index projections under future climate scenarios.

**Figure 2.**
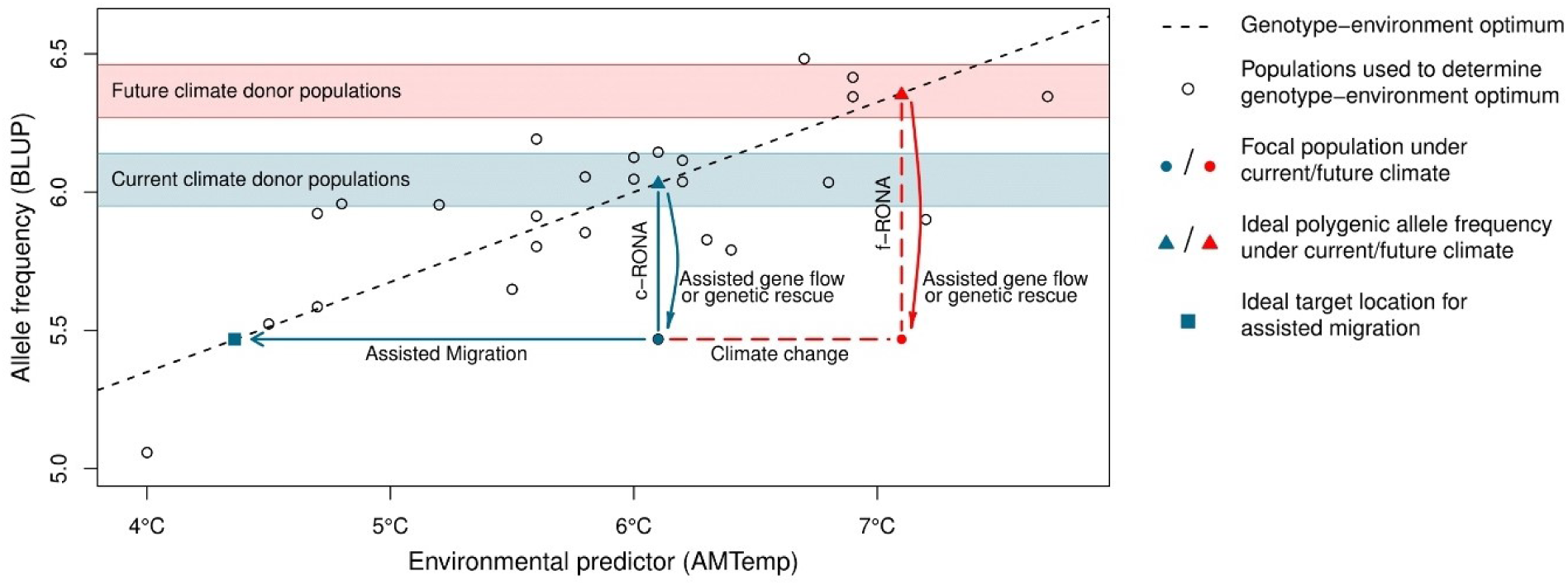
Schematic diagram of current and future risk of non-adaptedness (c-RONA and f-RONA), presented on a genotype-environment association (GEA) plot; where genotypes are BLUP estimates of population polygenic allele frequency for 17 loci and the environmental predictor is Annual Mean Temperature. c/f-RONA is the average change in allele frequency required to match our estimated optimum for current environmental conditions. Where RONA is large, we show two possible adaptation strategies; i) Assisted migration indicates the change in environmental conditions required for a population to match a genotype-environment optimum. This could take the form of a translocation of individuals to a location with a more suitable climate (e.g. a higher elevation). ii) Assisted Gene Flow (which in small populations is equivalent to Genetic Rescue) proposes movement of genetic material from a donor population with allele frequencies predicted to be better suited to the environmental conditions at the focal population. We show that the allele frequency change is likely to be larger under an example future climate scenario of 1°C warming. Blue and red bands indicate suitable candidate donor populations for assisted gene flow under current and future scenarios respectively.

### Conservation prioritization

We compared the magnitude of c-RONA across dwarf birch populations with the Shapley index (Haake et al., 2007). The Shapley index prioritizes populations based on evolutionary isolation and contribution to overall diversity based on pairwise differentiation. Several similar metrics are widely used for conservation management (Collen et al., 2011; Gumbs et al., 2018; Jetz et al., 2014). Here, we used the method outlined in Volkmann et al. (2014), which maximizes within-species genetic diversity using a network approach implemented in NeighborNet (Bryant and Moulton, 2004; Huson and Bryant, 2006). We used linear regression to test for a relationship between absolute c-RONA values and the Shapley index for neutral and adaptive loci.

### Simulated assisted gene flow

For each environmental variable, and for each population in the study, we identified the population most appropriate for AGF based on the match between the local environment and the sum of ***fβ***. Where several suitable populations were identified within the confidence interval of our regression, we selected the location geographically closest to the recipient population, since there could be local adaptation to undetected environmental variables (cf. Boshier *et al*. 2015).

### Method validation and ENM-GEA comparison

To validate our model we tested the hypothesis that higher c-RONA values would be associated with the reduced performance of fitness proxies. Therefore we tested for a correlation between population c-RONA values for each environmental variable or their interactions and the response of i) square root transformed catkin counts and ii) germination rate across study populations. Finally, we tested for a correlation between the relative importance of environmental variables identified in our ENM and the number of GEA loci associated with each variable.

## Results

### Environmental niche models

The dwarf birch ENM was well parameterized with high mean test AUC (0.946 ±0.008) and a low mean test omission rate (0.09, p<0.001) at a logistic threshold of occurrence of 0.193. Four variables together contributed >85% to the predictive model performance including annual mean temperature (34.9%) and maximum temperature of the warmest month (22.1%) (Table 1). The resulting model is highly concordant with qualitative field observations and inspection of variable curves showed biologically plausible responses (Figure S3). Future projections show significant declines across the species’ range with persistent populations restricted to areas of higher elevation (Figures 1, S4). Excluding other anthropogenic pressures, under the most severe scenario (RCP8.5, 2081-2100), suitable habitat may be reduced to ∼1% of the current extent (Table S2).

### Phenotypic data and habitat suitability

Phenotypic data means are reported in Table S3. Germination success was assayed in 190 individuals, and averaged 7.6% for both years with 6.1% 100-day survival (i.e. 80% of those that germinated) with substantial variation among populations (Table S4). A single large outlier individual (Emblehope) produced an exceptionally large number of catkins strongly biasing results, thus was excluded from subsequent analysis. Present time habitat suitability index (HSI) estimates for dwarf birch ranged from 0.0006 to 0.81 (Table 2), with substantial declines under all future scenarios (Figure S4). We found a significant non-linear positive relationship between HSI and mean population catkin count (F_1,26_=7.50, P=0.011) as well as HSI and the proportion of seeds that germinated (F_1,16_=9.52, P=0.007) (Figure 1).

### RAD Sequencing and genotype-environment associations

After quality control, RAD sequencing produced 173,460,998 reads, of which 79.1% aligned to the *B. nana* genome. Subsequently 73.2% of aligned reads mapped to a single unique position. Three samples were excluded due to low coverage. After filtering we retained 14,889 SNPs over 8,727 contigs. These contigs together cover approximately a third of the dwarf birch genome assembly. Bayescan identified 382 putative outlier SNPs with a relaxed false discovery rate of 0.2 which were excluded during the generation of the Bayenv2 null covariance matrix. Subsequent GEA analysis detected 267 highly significant locus-environment associations, encompassing 303 SNPs (Table S5), with a single SNP from each locus retained for subsequent analysis. The most frequent associations were between mean diurnal range and 71 loci, and annual mean temperature and 64 loci, whereas variables such as temperature seasonality and mean temperature of driest or wettest quarters had comparatively few associated loci. Just six loci were in common between Bayescan and Bayenv2 detection methods, and Bayescan candidate loci did not report significantly higher BF scores compared to the dataset as a whole. A comparison between bayenv2 and RDA found highly significant correlation (R_2_ = 071, F_1,6_ = 14.76, p = 0.008) between methods, in the number of genotype-environment associations identified for each environmental variable (Table S6, Figure S5) suggesting that both methods are identifying a similar genomic pattern of adaptation.

### Expression of putative adaptive loci

The 267 loci mapped to 185 unique scaffolds in our reference genome. Based on RNAseq data, 35 candidate regions showed evidence of gene expression in flower tissue (19%), 15 showed gene expression in leaf tissue (8%) and 13 showed gene expression in both (7%). In comparison to the overall SNP dataset, we found that both flower (X^2^=23.14, p<0.001) and leaf (X^2^=8.59, p=0.003) expressed sequences are significantly over-represented among putatively adaptive loci.

### Potential for adaptation and conservation prioritization

The c-RONA based on environmentally associated SNPs under present climate varied from 0.07 (SE ±0.06) at Glen Cannich, to 0.39 (±0.24) at Beinn Enaiglair on the Western periphery of the species range (Table 2, S7). BLUP estimates for all variables are presented in Figure S6. Under future climate scenarios mean population f-RONA was greater than c-RONA increasing from 0.22 (±0.10) to a maximum of 0.27 (±0.11) under scenario RCP8.5 (Table S8), with substantial variation across populations and projections. We found positive correlation between c-RONA and the Shapley Index for neutral genetic diversity (R_2_=0.2, F_1,24_=5.895, p=0.023), despite a number of outliers as shown by the low correlation coefficient, but no such pattern for putative adaptive genetic diversity (R_2_=0.00, F_1,24_=0.003, p=0.983) (Figure 3). The Shapley Index for neutral diversity also strongly favoured a small number of relict and range edge populations dominated by drift (e.g. BG, SA, see Borrell et al., 2018) whereas for adaptive diversity, the range of values was narrower suggested more even support across populations. Therefore, the Shapley Index and our metric for maladaptation (c-RONA) provide very different ranking for conservation value (Table 2). A consensus ranking of populations is provided in Table S9.

**Figure 3.**
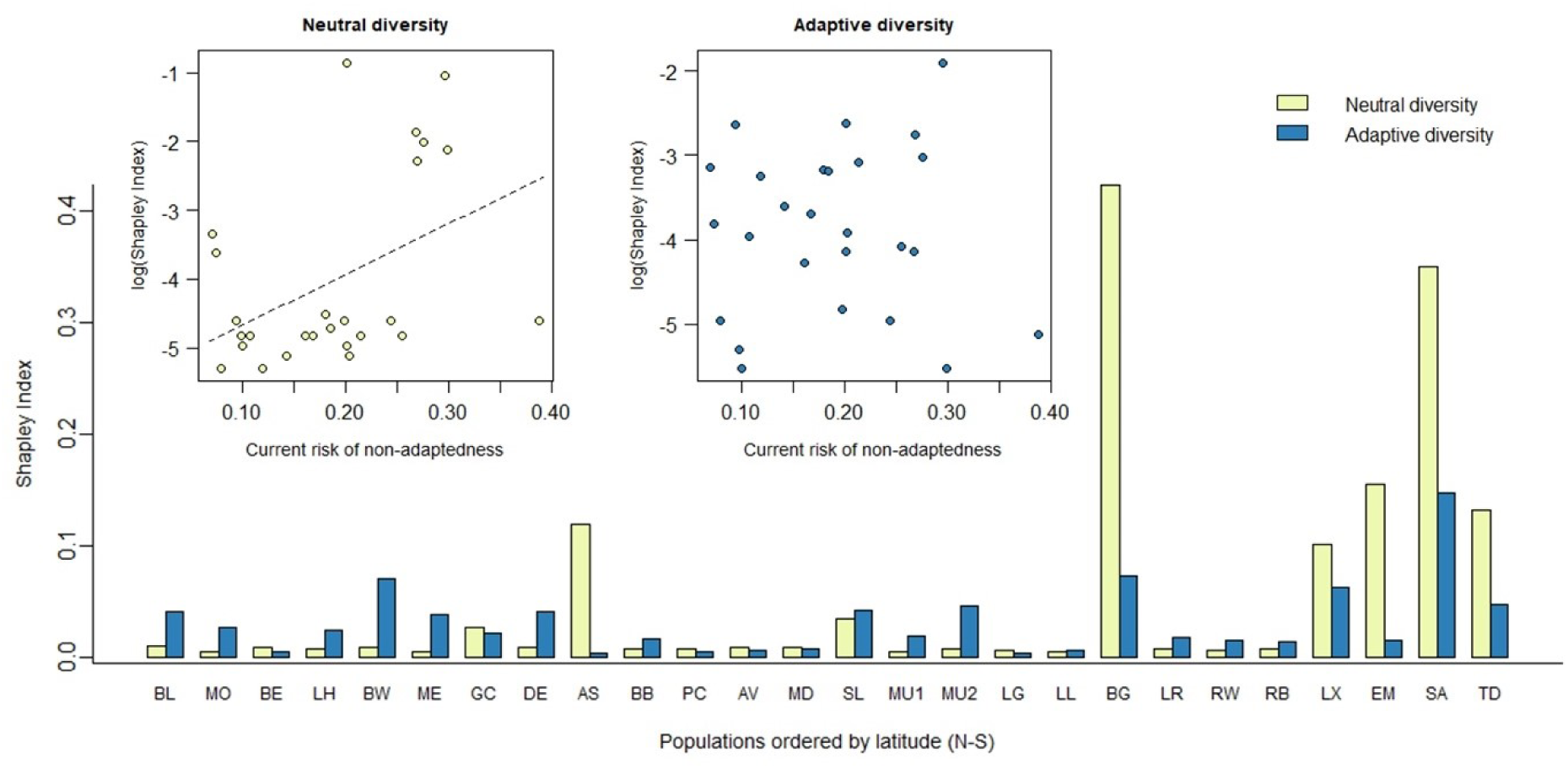
Barplot of Shapley index for neutral and adaptive loci across UK *B. nana* populations, ordered by latitude with northernmost populations to the left. Inset plots show the relationship between the log transformed Shapley Index and the current risk of non-adaptedness (c-RONA) for neutral and adaptive loci respectively.

### Simulating assisted gene flow

For each population across each environmental variable we identified the geographically closest ‘donor’ population with an allele frequency that would reduce c-RONA (within confidence limits) at the ‘recipient’ site (Figure 4, S7). This strategy proposes a pattern of dispersal from the centre of the distribution towards the periphery, particularly at the Southern range edge, though there are exceptions such as transfer from the Northern to Southern range edge (e.g. MTColdQ, Figure S7). In some cases, the analysis does not indicate the need for AGF in particular populations, such as those at the centre of the species distribution which appear to be well matched to their environment (i.e. locally adapted).

**Figure 4.**
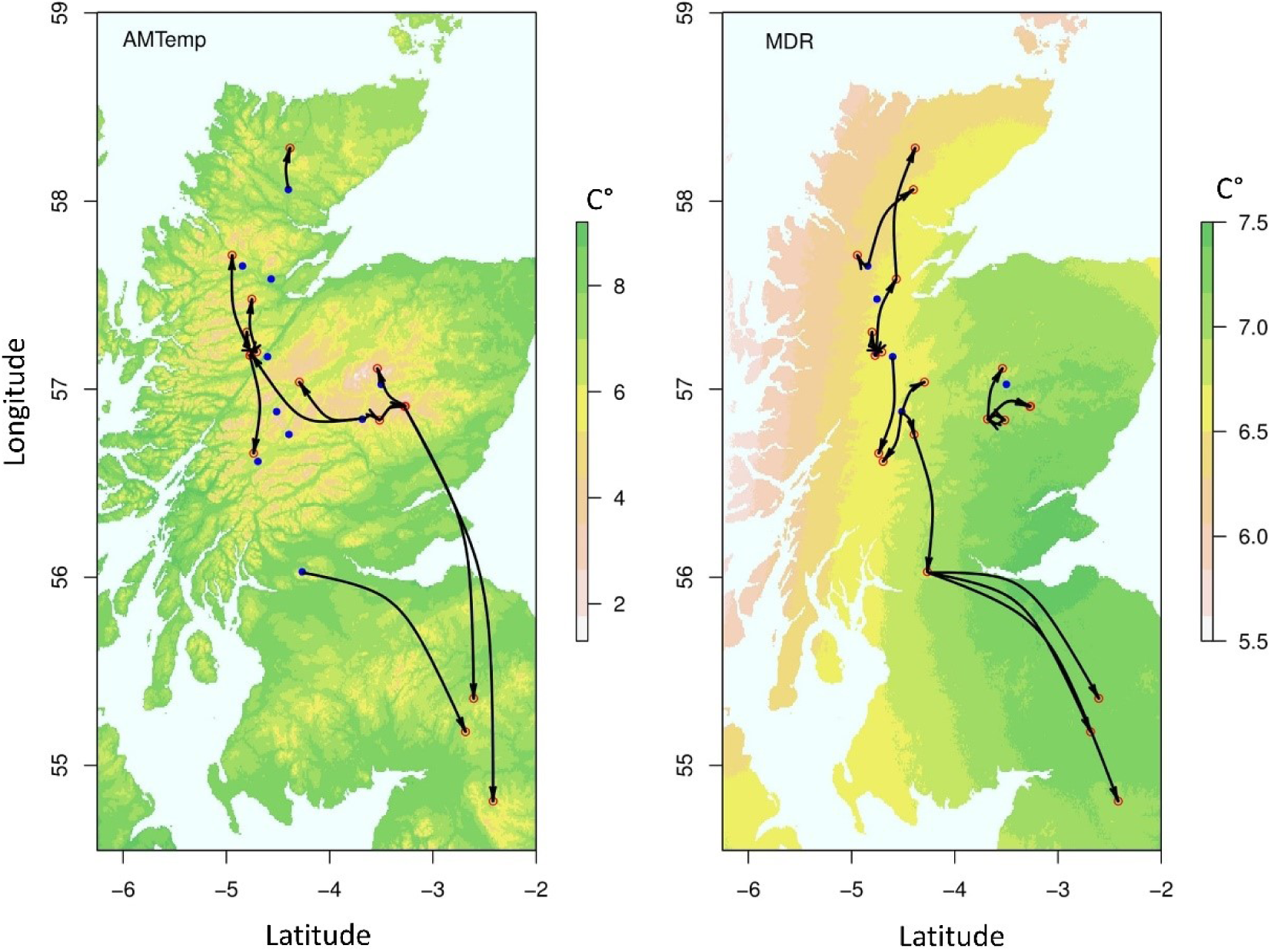
Hypothetical plots of assisted gene flow (AGF) for dwarf birch in the UK. Arrows denote movement from donor to recipient populations (red circles). Blue populations report an allele frequency close to predicted optimums, thus introduction of novel diversity does not decrease c-RONA and is not required. Base maps show Annual Mean Temperature (AMTemp) and Mean Diurnal Range (MDR) environmental variables.

### Method validation and ENM-GEA comparison

If c-RONA values do indeed quantify the degree of maladaptation, they should be negatively correlated with independent measurements of population fitness. The c-RONA values for annual mean temperature (AMTemp) were significantly negatively correlated with mean population catkin counts (F_1,23_=5.84, p=0.025) (Figure 5A) (we found a similar relationship for c-RONA averaged across all environmental variables, data not shown). The interaction of c-RONA for Annual Mean Temperature and Mean Diurnal Range correlated with germination rate (F_11,14_=8.07, p=0.004). Finally, in a comparison of ENM and GEA methods, we found a significant correlation between the number of genotype-environment associations and the percentage contribution of environmental variables defining species range in our ENM (F_1,8_ = 7.28, p = 0.027) (Figure 5B).

**Figure 5.**
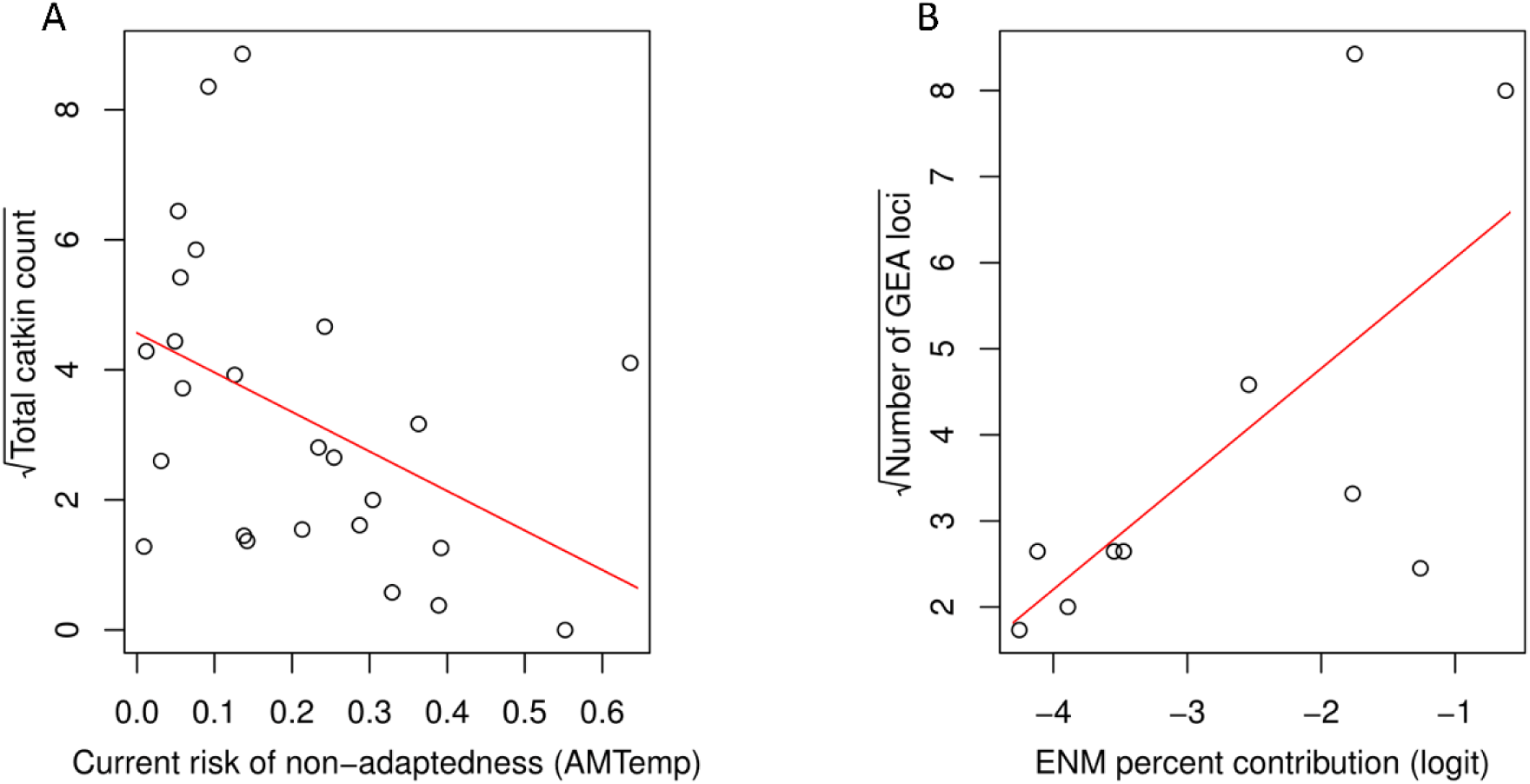
A) The relationship between c-RONA (for AMTemp) and mean population catkin count. B) Correlation between the number of loci identified in genotype-environment analyses, for each environmental variable, and the corresponding percentage contribution of that variable to the environmental niche model.

## Discussion

Environmental niche modelling projects that the decline of dwarf birch across the UK is likely to continue and become increasingly severe, with almost total range loss possible by the end of the century under the highest emission scenarios. We found that catkin production and seed germination are positively correlated with ENM projections of habitat suitability. This suggests lower reproductive fitness of plants in populations with lower habitat suitability index. We cannot fully exclude the possibility that low seed germination rates are partly due to high dormancy, but it is not obvious that dormancy would increase fitness unless it was a bet-hedging strategy for a plant in a poor environment. Temperature was particularly important to our ENM projections, and previous work has shown reduced production of germinable seeds by dwarf birch in warmer climates (Alsos *et al*. 2003). In future, an overall decline in habitat suitability across the species’ British range is likely to further reduce reproductive fitness and subsequent population persistence.

Genome-wide analysis identified 267 significant genotype-environment associations (0.018 of loci surveyed) across 24 environmental variables, which is consistent with the number of associations identified in similar studies (Abebe *et al*. 2015; Manthey and Moyle 2015; reviewed in Ahrens *et al*. 2018). These loci were significantly more commonly found within 10kb of a gene annotated on our reference genome sequence with cDNA evidence for expression than were SNP loci that were not identified as candidates, increasing our confidence that candidate loci could be involved in phenotypic traits.

We observe that of the four environmental variables that contribute substantially to the dwarf birch ENM (Table 1) three of these also account for the largest number of associated loci in the genotype-environment analysis (GEA) (Table 1, Table S5). Therefore, in a comparison of the two methods, we find significant agreement between ENM and GEA results in identifying important environmental variables (Figure 5B). It is not a logical necessity for environmental variables with the largest effects on species range limits to show the strongest correlation with allele frequencies. However, it is an interesting finding that suggests that we have identified biologically relevant environmental variables that influence both distribution and local adaptation of dwarf birch. It would be valuable to test for this pattern in other species, in the context of genetic models of species range limits (Polechová, 2018; Polechová and Barton, 2015).

We surveyed the allele frequencies of these GEA loci across populations to estimate c-RONA. As expected, we find the populations which we have identified as having a poor match between genotype and environment (high c-RONA) are particularly small or isolated, and those on the margins of the species’ niche. This result is consistent with reconstruction of demographic history and genetic differentiation by Borrell *et al*. (2018), who inferred that several of these small and isolated populations have been subject to severe genetic drift. We also found some of our c-RONA estimates or their interaction to correlate with catkin production and seed germination rates. This suggests low fitness due to maladaptation. We cannot exclude the possibility that reduced reproductive success could be an adaptive response to a poorer environment, but given the short timescales involved this seems unlikely.

Based on our inference that that populations with low c-RONA are more locally adapted, we then performed a comparison between c-RONA and the Shapley Index based on neutral diversity. We find that populations with the highest inferred conservation value (highest Shapley score for neutral loci) were also those with the greatest deviation from optimum allele frequencies (highest c-RONA) (Table 2, Figure 3). This implies that it may be inappropriate to use the Shapley Index (and by extension, other similar metrics) based solely on neutral diversity for conservation prioritization, since this strategy would inadvertently favour poorly adapted populations that display a high degree of unique variation – in the case of dwarf birch, this is most likely due to genetic drift. Instead, we propose a conservation framework where populations with a low c-RONA and high Shapley Index based instead on adaptive diversity are prioritized. This would maximize both local adaptation and adaptive diversity, supporting future adaptive potential (Table S9).

To illustrate a possible application for this prioritization framework, we sought to identify putative dwarf birch donor populations that possess adaptive alleles at frequencies that would display reduced c-RONA in a recipient population (Figures 4, S6). We chose to demonstrate our approach using a current climate reference, as it could be considered more conservative, though we note that planning for future climate may have a better chance of long-term success. In this example, our hypothetical AGF strategy involves a substantial translocation of genotypes, particularly from the centre of the range towards the periphery. Whilst controversial, AGF may be advantageous, as it can introduce or increase the frequency of preadapted alleles to allow more rapid adaptation to track changing climate, alleviate inbreeding depression or increase adaptive potential (Frankham, 2015; Prober et al., 2015); and in the process provide a demographic safeguard by augmenting population size (Hodgins and Moore, 2016). In practice, implementation of AGF is likely to take the form of composite provenancing, whereby genetic material from a combination of source populations is used (Breed et al., 2013; Hodgins and Moore, 2016). This may seek to target adaptive diversity across multiple important environmental variables from across the species range, sometimes irrespective of the distance to the source population and the ‘local is best’ paradigm (Boshier et al., 2015; Havens et al., 2015; Jones, 2013).

Our suggested approach has some limitations: RADseq only identifies variation in a subset of the genome (Lowry et al., 2016) possibly missing important adaptive loci (Harrisson et al., 2014). This concern may be addressed in future by whole genome population sequencing, and a better understanding of the limiting returns from typing more adaptive loci (for example Ahrens *et al*. 2018). Second, our approach does not explicitly account for phenotypic plasticity (which can be adaptive or non-adaptive), or the adaptive input from new mutations (Chevin and Lande, 2011). More generally, we caution against interpreting the statistical association between the RADseq alleles and the bioclimatic variates (for example, MDR) as a demonstration that the allele in question is linked to a quantitative trait locus with adaptive variation. Rather, the causal environmental variable may be unmeasured, but closely correlated with MDR. Finally, we highlight that, in our study area, the climate has been changing, albeit slowly, for several millennia, with the rate of climate change increasing more recently (Wang et al., 2014). Therefore, the clines identified here could represent adaptation to the environment of the recent past, rather than the present, and therefore may underestimate the current ecological risk. In the future, methods to accommodate change in the relative importance of environmental variables through time (Clark *et al*. 2014) and non-linear associations (Fitzpatrick and Keller 2015) are likely to advance our understanding and improve estimates of local adaptation in wild populations.

### Conclusions

Estimating the degree of maladaptation in populations as a criterion to inform selection of plant material for genetic rescue, composite provenancing or species reintroductions is currently the subject of considerable interest (Gibson et al., 2016; Leroy et al., 2018), and this is likely to increase in the context of environmental change (Aitken and Bemmels, 2016). Here we present an approach to permit rapid assessment of local adaptation and future adaptive potential in wild populations. Importantly, the estimation of maladaptation presents a testable hypothesis; specifically, that if an AGF programme translocated individuals to a site where they are expected to display reduced c-RONA, the response of measurable fitness proxies such as catkin production should be positive. In dwarf birch, AGF would have to be combined with other management interventions focused on mitigating burning and grazing pressure to support natural regeneration, with the aim that larger populations eventually support ‘natural’ gene flow. Similarly, AGF need not entail translocation of genetic material to an existing recipient population in the first instance. Initially individuals of different provenance (and known allele frequencies) could be translocated to trial locations and subsequent fitness assessments would enable validation of the predicted adaptive potential. Conservationists and practitioners would then be in a better position to manage and, where appropriate, facilitate adaptation.

## Acknowledgements

This work was funded by NERC Fellowship NE/G01504X/1 to Richard Buggs. James Borrell was funded by NERC CASE studentship NE/J017388/1 in collaboration with Trees for Life and Highland Birchwoods. Jasmin Zohren was funded by the Marie-Curie Initial Training Network INTERCROSSING. We gratefully acknowledge helpful comments from Dr. Chris Eizaguirre and the assistance provided by members of the Montane Scrub Action Group and numerous landowners who permitted access to their estates.

## Data Archiving Statement

1. Illumina read data from RADseq libraries has been uploaded to the European Nucleotide Archive project PRJEB26807, sample accessions ERS2598190-ERS2598376
2. Species records are available directly from the NBN Gateway [https://data.nbn.org.uk/].
3. Climate data are available from http://www.worldclim.org/

## Supplementary Materials

### Phenotyping and germination protocol

All UK populations were visited once or twice in the spring and summer of 2012, 2013 or 2014, once plants were in leaf to aid identification. For each individual, the following phenotypic measurements were made:

- Latitude and longitude (GPS: Garmin Oregon 550)
- Elevation (GPS based)
- Number of male and female catkins.
- Perpendicular eight from ground level.
- Browsing pressure (percentage of browsed stems, to nearest 5%).
- Plant area (length of the longest horizontal growing axis multiplied by maximum width perpendicular to this)
- Diameter of the largest available stem at ground level.

In the years 2013 and 2014, seeds were collected from a subset of 18 populations (9 per year) in Scotland to assess germination rates. We ensured that collected catkins displayed dry brown bracts and readily dehisced to ensure maturity. Catkins were placed in labelled glassine envelopes and further air-dried for 3-5 days before being stored at 4°C for planting the following Spring. It should be noted that collected catkins would have been from the previous year’s growth, so not necessarily correlated to the female catkin count also reported in this study.

To assay germination, seeds were counted and spread on filter paper in individually labelled petri dishes. Where a large amount of seed was available for a given individual, petri dishes were replicated to avoid overcrowding. A thin layer of vermiculite was then added to prevent desiccation. Seeds were maintained at 18-20°C with a 14h photoperiod for 60 days. Germination was scored twice weekly and considered successful where a radicle ≥ 5mm was observed. For populations assayed in 2014, successfully germinated seedlings were transferred to a nutrient poor soil (similar to their preferred habitat) to assess survivability at 100 days.

### Redundancy Analysis of genotype-environment associations

For comparison, we also tested the pattern of genotype-environment associations (GEA) using Redundancy Analysis (RDA), a method that has shown robust performance in scenarios of weak selection (Forester et al., 2018; Rellstab et al., 2015). RDA is a two-step analysis which extends multivariate linear regression to allow regression of multiple response variables on multiple explanatory variables. A PCA of the fitted values results in canonical axes which are linear combinations of the environmental predictors, therefore permitting identification of significant GEAs (Legendre and Legendre, 2012).

We implemented RDA in the R package vegan (Oksanen et al., 2019), using the full 14,889 SNP dataset, and a reduced set of environmental variables. Whilst we used all environmental predictor variables across independent runs of the Bayenv2 analysis (main text), here we use a reduced set of environmental variables to avoid correlated predictors being analyzed together. The reduced set of environmental variables was the same as those used for environmental niche modelling (n=10), with the additional exclusion of MTDryQ and MTWet (n=8), which showed collinearity > 0.7 in this reduced number of population sampling locations (variables for the ENM were assessed across the whole study area).

We followed the methodology outlined in (Forester et al., 2018), retaining candidate SNPs from the first three axes, with a 2.5 standard deviation significance threshold. For each candidate SNP, we first identified the environmental predictor with which it reported the highest correlation. Second, we compare candidates to those identified in the Bayenv2 GEA analysis (main text). Finally, we compare the number of SNPs associated with each environmental predictor variable across both RDA and Bayenv2 methods.

RDA identified 601 significant genotype-environment associations across eight retained predictor variables (Table S6). In a comparison of candidates between GEA methods, 11.2% of significant Bayenv2 loci were also significant in the RDA analysis. This is consistent with 9.4% of loci found in common between Bayenv2 and RDA analyses in Schweizer *et al*. (2016) and Forester *et al*. (2018). Finally, we report a highly significant correlation between the number of associations identified for each environmental variable using RDA and Bayenv2 (F_1,6_ = 14.76, p = 0.008), Figure S5). We note that this pattern was significant across a range of RDA significance thresholds, as well as with both the loci directly associated with retained variables, and the loci correlated with retained variables (see columns 5 and 6, Table 1), therefore we are satisfied it is a robust and repeatable pattern.

## Supplementary Tables

**Table S1.** Projected change in global mean surface air temperature, relative to the period 1986–2005. Adapted from (IPCC, 2014b).

| Scenario | 2046-2065 | 2081-2100 |
| --- | --- | --- |
| | Mean $\Delta$ °C (Likely range) | Mean $\Delta$ °C (Likely range) |
| RCP2.6 | +1.0 (0.4 to 1.6) | +1.0 (0.3 to 1.7) |
| RCP4.5 | +1.4 (0.9 to 2.0) | +1.8 (1.1 to 2.6) |
| RCP6.0 | +1.3 (0.8 to 1.8) | +2.2 (1.4 to 3.1) |
| RCP8.5 | +2.0 (1.4 to 2.6) | +3.7 (2.6 to 4.8) |

**Table S2.**
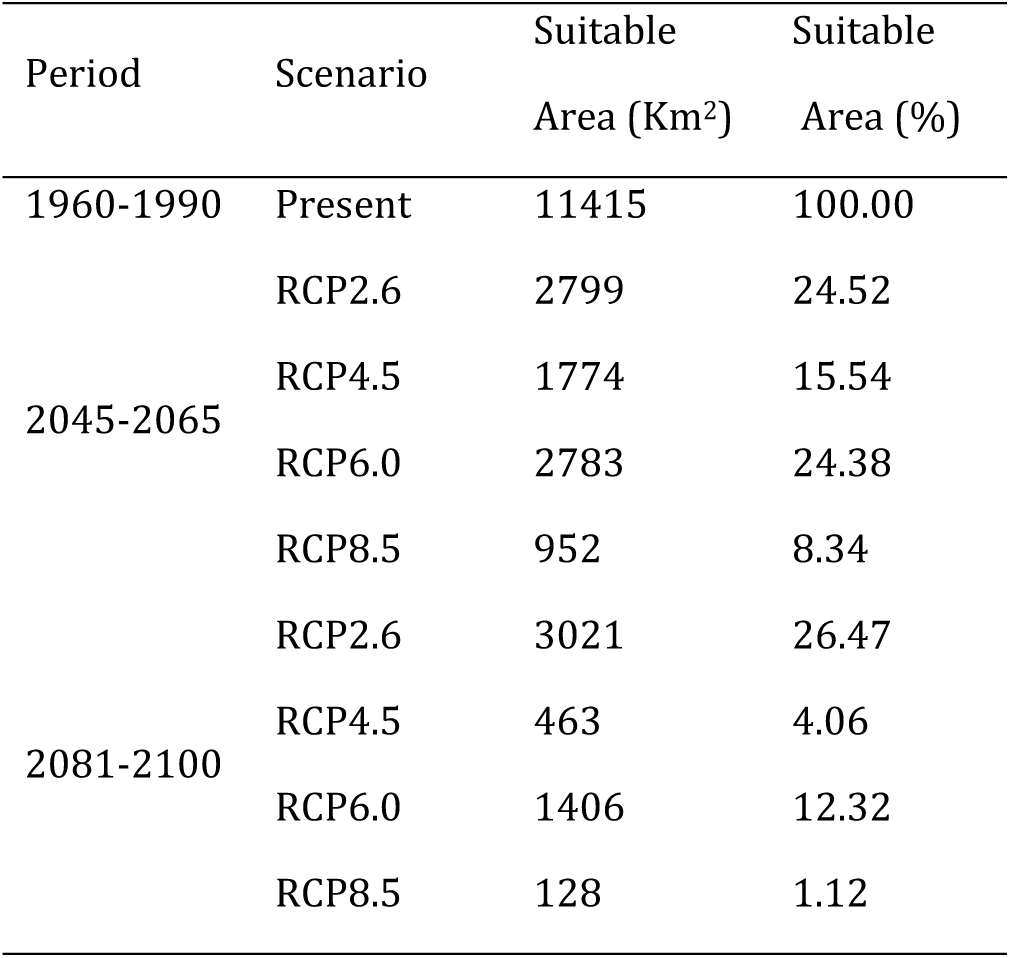
Changes in suitable habitat area as defined by ‘maximum training sensitivity plus specificity’ threshold for dwarf birch under IPCC future climate scenarios.

**Table S3.**
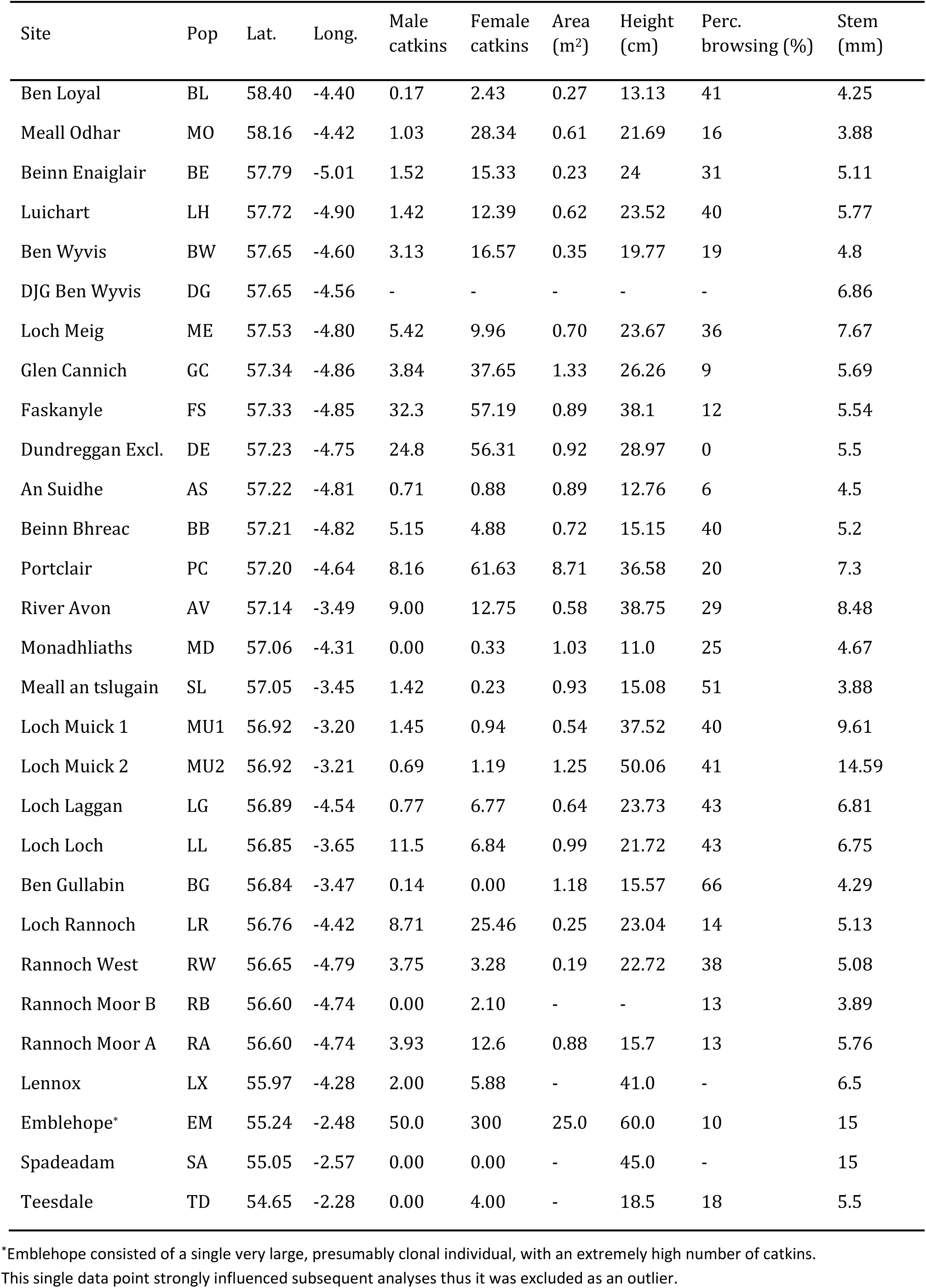
Phenotypic data summary for sampled dwarf birch populations.

| Site | Pop | Lat. | Long. | Male catkins | Female catkins | Area (m<sup>2</sup>) | Height (cm) | Perc. browsing (%) | Stem (mm) |
| --- | --- | --- | --- | --- | --- | --- | --- | --- | --- |
| Ben Loyal | BL | 58.40 | -4.40 | 0.17 | 2.43 | 0.27 | 13.13 | 41 | 4.25 |
| Meall Odhar | MO | 58.16 | -4.42 | 1.03 | 28.34 | 0.61 | 21.69 | 16 | 3.88 |
| Beinn Enaiglair | BE | 57.79 | -5.01 | 1.52 | 15.33 | 0.23 | 24 | 31 | 5.11 |
| Luichart | LH | 57.72 | -4.90 | 1.42 | 12.39 | 0.62 | 23.52 | 40 | 5.77 |
| Ben Wyvis | BW | 57.65 | -4.60 | 3.13 | 16.57 | 0.35 | 19.77 | 19 | 4.8 |
| DJG Ben Wyvis | DG | 57.65 | -4.56 | - | - | - | - | - | 6.86 |
| Loch Meig | ME | 57.53 | -4.80 | 5.42 | 9.96 | 0.70 | 23.67 | 36 | 7.67 |
| Glen Cannich | GC | 57.34 | -4.86 | 3.84 | 37.65 | 1.33 | 26.26 | 9 | 5.69 |
| Faskanyle | FS | 57.33 | -4.85 | 32.3 | 57.19 | 0.89 | 38.1 | 12 | 5.54 |
| Dundreggan Excl. | DE | 57.23 | -4.75 | 24.8 | 56.31 | 0.92 | 28.97 | 0 | 5.5 |
| An Suidhe | AS | 57.22 | -4.81 | 0.71 | 0.88 | 0.89 | 12.76 | 6 | 4.5 |
| Beinn Bhreac | BB | 57.21 | -4.82 | 5.15 | 4.88 | 0.72 | 15.15 | 40 | 5.2 |
| Portclair | PC | 57.20 | -4.64 | 8.16 | 61.63 | 8.71 | 36.58 | 20 | 7.3 |
| River Avon | AV | 57.14 | -3.49 | 9.00 | 12.75 | 0.58 | 38.75 | 29 | 8.48 |
| Monadhliaths | MD | 57.06 | -4.31 | 0.00 | 0.33 | 1.03 | 11.0 | 25 | 4.67 |
| Meall an tslugain | SL | 57.05 | -3.45 | 1.42 | 0.23 | 0.93 | 15.08 | 51 | 3.88 |
| Loch Muick 1 | MU1 | 56.92 | -3.20 | 1.45 | 0.94 | 0.54 | 37.52 | 40 | 9.61 |
| Loch Muick 2 | MU2 | 56.92 | -3.21 | 0.69 | 1.19 | 1.25 | 50.06 | 41 | 14.59 |
| Loch Laggan | LG | 56.89 | -4.54 | 0.77 | 6.77 | 0.64 | 23.73 | 43 | 6.81 |
| Loch Loch | LL | 56.85 | -3.65 | 11.5 | 6.84 | 0.99 | 21.72 | 43 | 6.75 |
| Ben Gullabin | BG | 56.84 | -3.47 | 0.14 | 0.00 | 1.18 | 15.57 | 66 | 4.29 |
| Loch Rannoch | LR | 56.76 | -4.42 | 8.71 | 25.46 | 0.25 | 23.04 | 14 | 5.13 |
| Rannoch West | RW | 56.65 | -4.79 | 3.75 | 3.28 | 0.19 | 22.72 | 38 | 5.08 |
| Rannoch Moor B | RB | 56.60 | -4.74 | 0.00 | 2.10 | - | - | 13 | 3.89 |
| Rannoch Moor A | RA | 56.60 | -4.74 | 3.93 | 12.6 | 0.88 | 15.7 | 13 | 5.76 |
| Lennox | LX | 55.97 | -4.28 | 2.00 | 5.88 | - | 41.0 | - | 6.5 |
| Emblehope* | EM | 55.24 | -2.48 | 50.0 | 300 | 25.0 | 60.0 | 10 | 15 |
| Spadeadam | SA | 55.05 | -2.57 | 0.00 | 0.00 | - | 45.0 | - | 15 |
| Teesdale | TD | 54.65 | -2.28 | 0.00 | 4.00 | - | 18.5 | 18 | 5.5 |

\*Emblehope consisted of a single very large, presumably clonal individual, with an extremely high number of catkins. This single data point strongly influenced subsequent analyses thus it was excluded as an outlier.

**Table S4.** Germination success and survivability summary data for assayed populations.

| Year | Population | Individuals | Seeds<br>Planted | Germinated | Germ. % | 100-Day<br>Survivability | Surv. % |
| --- | --- | --- | --- | --- | --- | --- | --- |
| 2013 | AV | 13 | 438 | 1 | 0.23 | - | - |
| 2013 | BB | 7 | 102 | 2 | 1.96 | - | - |
| 2013 | BL | 2 | 35 | 0 | 0.00 | - | - |
| 2013 | DE | 17 | 540 | 24 | 4.44 | - | - |
| 2013 | GC | 8 | 833 | 68 | 8.16 | - | - |
| 2013 | LL | 10 | 187 | 0 | 0.00 | - | - |
| 2013 | LR | 6 | 63 | 0 | 0.00 | - | - |
| 2013 | ME | 3 | 67 | 0 | 0.00 | - | - |
| 2013 | MU | 8 | 151 | 0 | 0.00 | - | - |
| Total<br>2013 |  | 74 | 2416 | 95 | 3.93 | - | - |
| 2014 | DJG | 21 | 1345 | 134 | 9.96 | 27 | 2.01 |
| 2014 | FS | 23 | 492 | 89 | 18.09 | 86 | 17.48 |
| 2014 | LG | 31 | 310 | 16 | 5.16 | 15 | 4.84 |
| 2014 | LX | 5 | 230 | 0 | 0.00 | 0 | 0.00 |
| 2014 | RA | 2 | 31 | 1 | 3.23 | 0 | 0.00 |
| 2014 | RB | 3 | 21 | 0 | 0.00 | 0 | 0.00 |
| 2014 | PC | 28 | 672 | 101 | 15.03 | 77 | 11.46 |
| 2014 | TD | 2 | 14 | 0 | 0.00 | 0 | 0.00 |
| 2014 | EM | 1 | 250 | 5 | 2.00 | 1 | 0.40 |
| Total<br>2014 |  | 116 | 3365 | 346 | 10.28 | 206 | 6.12 |

**Table S5.** The 24 environmental variables included in this study. Uncorrelated retained environmental variables were used for Environmental Niche Modelling (ENM), whilst all variables were tested independently for genotype-environment associations (GEA).

| Variable | Description | Retained<br>for ENM | Grouping | Bayenv2<br>GEA Loci<br>(totals inc. cor.) |
| --- | --- | --- | --- | --- |
| AMTemp | Annual Mean Temperature | X | A | 17 (64) |
| MTColdQ | Mean Temperature of Coldest Quarter |  | A | 24 |
| MTColdM | Min Temperature of Coldest Month |  | A | 23 |
| MTWarmM | Max Temperature of Warmest Month | X | B | 2 (6) |
| MTWarmQ | Mean Temperature of Warmest Quarter |  | B | 4 |
| MDR | Mean Diurnal Temperature Range | X | C | 71 (71) |
| ISO | Isothermality | X | D | 11 (11) |
| APrec | Annual Precipitation | X | E | 2 (21) |
| PWetQ | Precipitation of Wettest Quarter |  | E | 2 |
| PDryQ | Precipitation of Driest Quarter |  | E | 4 |
| PWetM | Precipitation of Wettest Month |  | E | 2 |
| PDryM | Precipitation of Driest Month |  | E | 3 |
| Pseason | Precipitation Seasonality |  | E | 1 |
| PWarmQ | Precipitation of Warmest Quarter |  | E | 4 |
| PColdQ | Precipitation of Coldest Quarter |  | E | 3 |
| Slope | Slope (derived from elevation) | X | F | 7 (7) |
| MTDryQ | Mean Temperature of Driest Quarter | X | G | 7 (7) |
| Tseason | Temperature Seasonality | X | H | 1 (3) |
| ATempR | Annual Temperature Range |  | H | 2 |
| MTWetQ | Mean Temperature of Wettest Quarter | X | I | 7 (7) |
| Aspect | Aspect (derived from elevation) | X | J | 4 (4) |
| Elev. | Elevation | - | - | 12 |
| Lat. | Latitude | - | - | 6 |
| Long. | Longitude | - | - | 48 |

**Table S6.** Comparison of GEA candidate loci identified in RDA and Bayenv2 analysis

| Variable | GEA Loci | GEA Loci (inc. cor.) | RDA (s.d. = 3) | RDA (s.d. =2.5 |
| --- | --- | --- | --- | --- |
| AMTemp | 17 | 64 | 21 | 101 |
| MTWarmM | 2 | 6 | 3 | 37 |
| MDR | 71 | 71 | 40 | 134 |
| ISO | 11 | 11 | 16 | 69 |
| APrec | 2 | 21 | 20 | 82 |
| Slope | 7 | 7 | 17 | 93 |
| MTDryQ | 7 | 7 | - | - |
| TS | 1 | 3 | 13 | 41 |
| MTWetQ | 7 | 7 | - | - |
| Aspect | 4 | 4 | 13 | 44 |

**Table S7.** Current risk of non-adaptedness across all genotype-environment analyses for retained environmental variables.

| pop | AMTemp | MDR | ISO | MTColdM | MTWetQ | MTDryQ | MTColdQ | Slope | Elev. | Combined |
| --- | --- | --- | --- | --- | --- | --- | --- | --- | --- | --- |
| BL | 0.287 | 0.188 | 0.216 | 0.374 | 0.349 | 0.021 | 0.079 | 0.091 | 0.011 | 0.18 |
| MO | 0.056 | 0.232 | 0.237 | 0.117 | 0.167 | 0.251 | 0.156 | 0.043 | 0.018 | 0.14 |
| BE | 0.636 | 0.483 | 0.46 | 0.672 | 0.135 | 0.222 | 0.637 | 0.23 | 0.015 | 0.39 |
| LH | 0.059 | 0.069 | 0.48 | 0.307 | 0.009 | 0.39 | 0.057 | 0.131 | 0.006 | 0.17 |
| BW | 0.049 | 0.283 | 0.318 | 0.017 | 0.02 | 0.08 | 0.028 | 0.05 | 0.003 | 0.09 |
| ME | 0.126 | 0.135 | 0.081 | 0.078 | 0.013 | 0.383 | 0.22 | 0.015 | 0.018 | 0.12 |
| GC | 0.053 | 0.017 | 0.077 | 0.082 | 0.064 | 0.196 | 0.006 | 0.157 | 0.013 | 0.07 |
| DE | 0.136 | 0.159 | 0.353 | 0.048 | 0.114 | 0.373 | 0.323 | 0.144 | 0.012 | 0.18 |
| AS | 0.392 | 0.085 | 0.144 | 0.347 | 0.02 | 0.559 | 0.267 | 0.843 | 0.032 | 0.30 |
| BB | 0.363 | 0.469 | 0.3 | 0.337 | 0.056 | 0.146 | 0.516 | 0.106 | 0.005 | 0.26 |
| PC | 0.092 | 0.058 | 0.071 | 0.166 | 0.044 | 0.297 | 0.036 | 0.115 | 0.001 | 0.10 |
| AV | 0.242 | 0.317 | 0.348 | 0.464 | 0.014 | 0.268 | 0.443 | 0.08 | 0.016 | 0.24 |
| MD | 0.329 | 0.125 | 0.078 | 0.377 | 0.061 | 0.09 | 0.554 | 0.162 | 0.006 | 0.20 |
| SL | 0.009 | 0.117 | 0.111 | 0.09 | 0.033 | 0.102 | 0.067 | 0.065 | 0.032 | 0.07 |
| MU1 | 0.213 | 0.311 | 0.561 | 0.09 | 0.012 | 0.536 | 0.066 | 0.027 | 0.009 | 0.20 |
| MU2 | 0.142 | 0.314 | 0.55 | 0.036 | 0.089 | 0.161 | 0.063 | 0.566 | 0.003 | 0.21 |
| LG | 0.031 | 0.05 | 0.164 | 0.031 | 0.076 | 0.292 | 0.027 | 0.226 | 0.001 | 0.10 |
| LL | 0.012 | 0.149 | 0.23 | 0.026 | 0.005 | 0.026 | 0.161 | 0.075 | 0.025 | 0.08 |
| BG | 0.389 | 0.204 | 0.05 | 0.041 | 0.063 | 0.769 | 0.23 | 0.039 | 0.026 | 0.20 |
| LR | 0.076 | 0.125 | 0.141 | 0.051 | 0.003 | 0.53 | 0.02 | 0.013 | 0.001 | 0.11 |
| RW | 0.254 | 0.258 | 0.109 | 0.309 | 0.2 | 0.509 | 0.099 | 0.072 | 0.002 | 0.20 |
| RB | 0.138 | 0.16 | 0.402 | 0.253 | 0.063 | 0.155 | 0.159 | 0.112 | 0.007 | 0.16 |
| LX | 0.234 | 0.17 | 0.561 | 0.22 | 0.325 | 0.455 | 0.41 | 0.046 | 0.001 | 0.27 |
| EM | 0.383 | 0.292 | 0.479 | 0.128 | 0.175 | 0.756 | 0.109 | 0.084 | 0.008 | 0.27 |
| SA | 0.552 | 0.317 | 0.54 | 0.541 | 0.023 | 0.464 | 0.109 | 0.107 | 0.011 | 0.30 |
| TD | 0.304 | 0.211 | 0.149 | 0.558 | 0.174 | 0.55 | 0.491 | 0.038 | 0.008 | 0.28 |

**Table S8.** Risk of non-adaptedness under current and future climate scenarios, excluding associations with altitude and slope.

| Pop | c-RONA<br>(no-elev/slope) | f-RONA (2045-2065) |  |  |  | f-RONA (2081-2100) |  |  |  |
| --- | --- | --- | --- | --- | --- | --- | --- | --- | --- |
|  |  | RCP2.6 | RCP4.5 | RCP6.0 | RCP8.5 | RCP2.6 | RCP4.5 | RCP6.0 | RCP8.5 |
| BL | 0.216 | 0.272 | 0.311 | 0.262 | 0.375 | 0.280 | 0.236 | 0.254 | 0.242 |
| MO | 0.174 | 0.243 | 0.286 | 0.231 | 0.360 | 0.247 | 0.247 | 0.255 | 0.218 |
| BE | 0.463 | 0.455 | 0.478 | 0.458 | 0.521 | 0.454 | 0.405 | 0.483 | 0.426 |
| LH | 0.196 | 0.167 | 0.243 | 0.190 | 0.306 | 0.187 | 0.199 | 0.169 | 0.189 |
| BW | 0.113 | 0.135 | 0.205 | 0.104 | 0.218 | 0.127 | 0.089 | 0.134 | 0.103 |
| ME | 0.148 | 0.255 | 0.104 | 0.244 | 0.165 | 0.225 | 0.203 | 0.233 | 0.202 |
| GC | 0.071 | 0.182 | 0.076 | 0.159 | 0.086 | 0.188 | 0.116 | 0.188 | 0.113 |
| DE | 0.215 | 0.338 | 0.275 | 0.175 | 0.274 | 0.338 | 0.231 | 0.304 | 0.235 |
| AS | 0.259 | 0.278 | 0.220 | 0.274 | 0.223 | 0.294 | 0.267 | 0.300 | 0.305 |
| BB | 0.312 | 0.398 | 0.392 | 0.433 | 0.354 | 0.417 | 0.294 | 0.419 | 0.323 |
| PC | 0.109 | 0.117 | 0.202 | 0.107 | 0.163 | 0.114 | 0.157 | 0.112 | 0.161 |
| AV | 0.299 | 0.333 | 0.407 | 0.296 | 0.323 | 0.322 | 0.300 | 0.317 | 0.290 |
| MD | 0.230 | 0.422 | 0.374 | 0.453 | 0.344 | 0.415 | 0.214 | 0.436 | 0.226 |
| SL | 0.075 | 0.080 | 0.108 | 0.085 | 0.186 | 0.074 | 0.048 | 0.081 | 0.088 |
| MU1 | 0.255 | 0.244 | 0.210 | 0.243 | 0.320 | 0.236 | 0.228 | 0.258 | 0.240 |
| MU2 | 0.193 | 0.241 | 0.261 | 0.206 | 0.310 | 0.207 | 0.211 | 0.233 | 0.205 |
| LG | 0.096 | 0.160 | 0.136 | 0.180 | 0.135 | 0.170 | 0.123 | 0.167 | 0.133 |
| LL | 0.087 | 0.078 | 0.219 | 0.087 | 0.201 | 0.089 | 0.081 | 0.069 | 0.086 |
| BG | 0.250 | 0.254 | 0.185 | 0.251 | 0.124 | 0.237 | 0.240 | 0.238 | 0.232 |
| LR | 0.135 | 0.130 | 0.083 | 0.123 | 0.075 | 0.142 | 0.095 | 0.093 | 0.104 |
| RW | 0.248 | 0.240 | 0.236 | 0.206 | 0.243 | 0.170 | 0.212 | 0.236 | 0.263 |
| RB | 0.190 | 0.159 | 0.194 | 0.184 | 0.234 | 0.190 | 0.216 | 0.203 | 0.172 |
| LX | 0.339 | 0.523 | 0.538 | 0.497 | 0.387 | 0.535 | 0.327 | 0.523 | 0.350 |
| EM | 0.332 | 0.387 | 0.398 | 0.397 | 0.388 | 0.260 | 0.324 | 0.242 | 0.369 |
| SA | 0.364 | 0.329 | 0.360 | 0.322 | 0.333 | 0.357 | 0.374 | 0.344 | 0.375 |
| TD | 0.348 | 0.421 | 0.385 | 0.433 | 0.365 | 0.431 | 0.355 | 0.436 | 0.342 |
| Mean | 0.220 | 0.263 | 0.265 | 0.254 | 0.270 | 0.258 | 0.223 | 0.259 | 0.230 |

**Table S9.** Shapley values for neutral and putative adaptive loci, and population c-RONA ordered by rank. Final column represents a consensus ranking with Shapley Index for adaptive loci maximized and c-RONA minimized to optimize both adaptive diversity and current local adaptation.

| Pop | Shapley (neutral) | Pop | Shapley (adaptive) | Pop | c-RONA | Consensus rank |
| --- | --- | --- | --- | --- | --- | --- |
| BG | 0.422 | SA | 0.148 | GC | 0.045 | GC |
| SA | 0.350 | BG | 0.073 | LG | 0.064 | SL |
| EM | 0.155 | BW | 0.071 | PC | 0.081 | BW |
| TD | 0.133 | LX | 0.063 | SL | 0.085 | LR |
| AS | 0.119 | TD | 0.048 | LR | 0.097 | BL |
| LX | 0.102 | MU2 | 0.046 | LL | 0.106 | DE |
| SL | 0.035 | SL | 0.043 | ME | 0.128 | ME |
| GC | 0.027 | BL | 0.042 | LH | 0.131 | LH |
| BL | 0.011 | DE | 0.041 | BW | 0.149 | MO |
| AV | 0.010 | ME | 0.039 | MO | 0.168 | BG |
| BW | 0.010 | MO | 0.027 | RB | 0.169 | RB |
| MD | 0.010 | LH | 0.025 | DE | 0.174 | LX |
| BE | 0.010 | GC | 0.022 | BL | 0.194 | MU2 |
| DE | 0.009 | MU1 | 0.020 | BG | 0.194 | MU1 |
| LR | 0.008 | LR | 0.019 | MU2 | 0.218 | RW |
| BB | 0.008 | BB | 0.017 | RW | 0.218 | TD |
| RB | 0.008 | RW | 0.016 | AS | 0.219 | LL |
| PC | 0.008 | EM | 0.016 | MD | 0.222 | MD |
| MU2 | 0.008 | RB | 0.014 | MU1 | 0.223 | EM |
| LH | 0.008 | MD | 0.008 | LX | 0.241 | BB |
| RW | 0.007 | AV | 0.007 | EM | 0.254 | AV |
| LG | 0.007 | LL | 0.007 | TD | 0.291 | PC |
| MO | 0.006 | BE | 0.006 | AV | 0.306 | SA |
| MU1 | 0.006 | PC | 0.005 | SA | 0.321 | LG |
| ME | 0.005 | LG | 0.004 | BB | 0.366 | AS |
| LL | 0.005 | AS | 0.004 | BE | 0.479 | BE |

## Supplementary Figures

**Figure S1.**
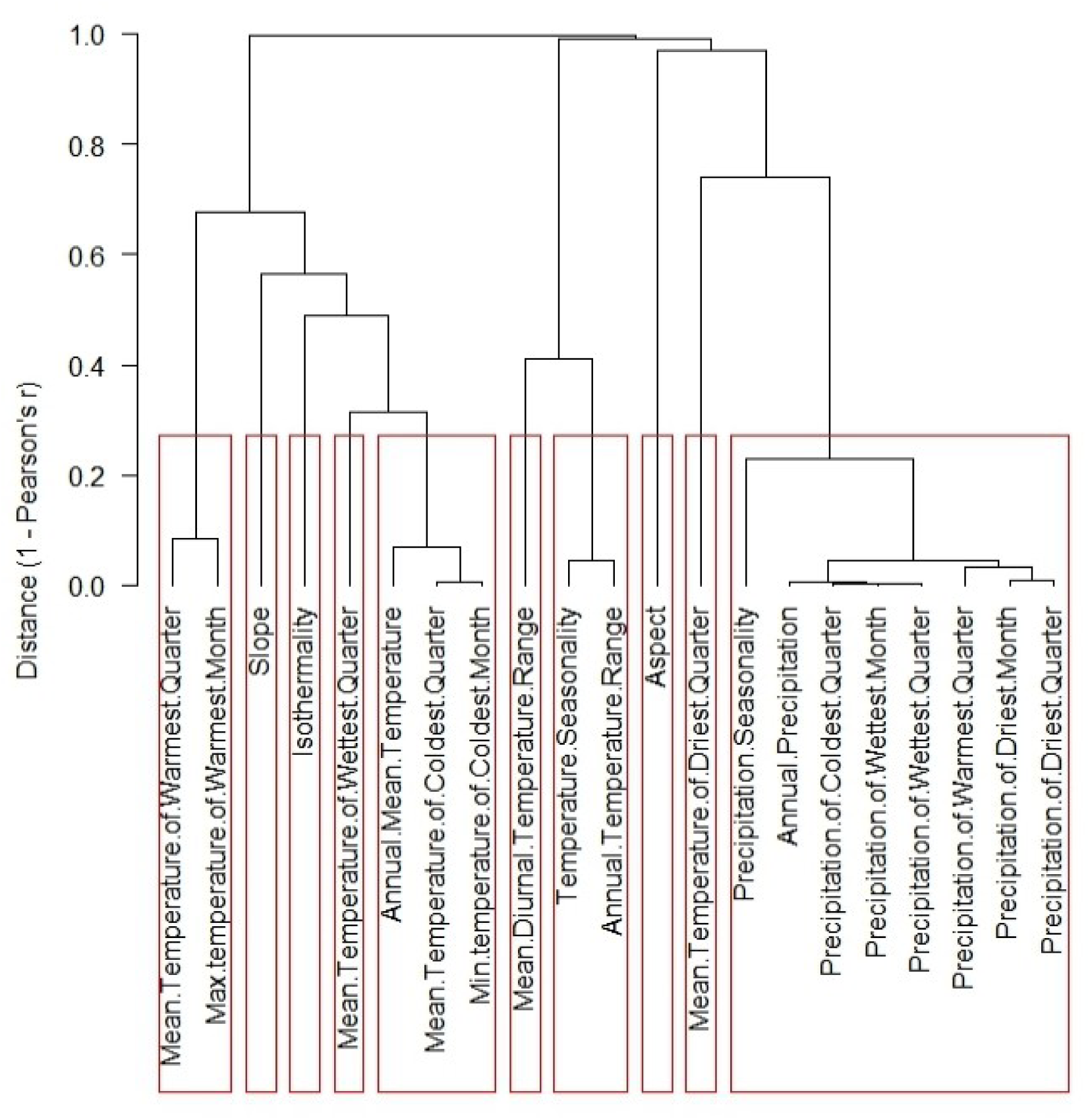
Topology of collinearity between environmental variables used in this study, at a threshold of 0.7. Red boxes denote groups of retained variables (see Table S5).

**Figure S2.**
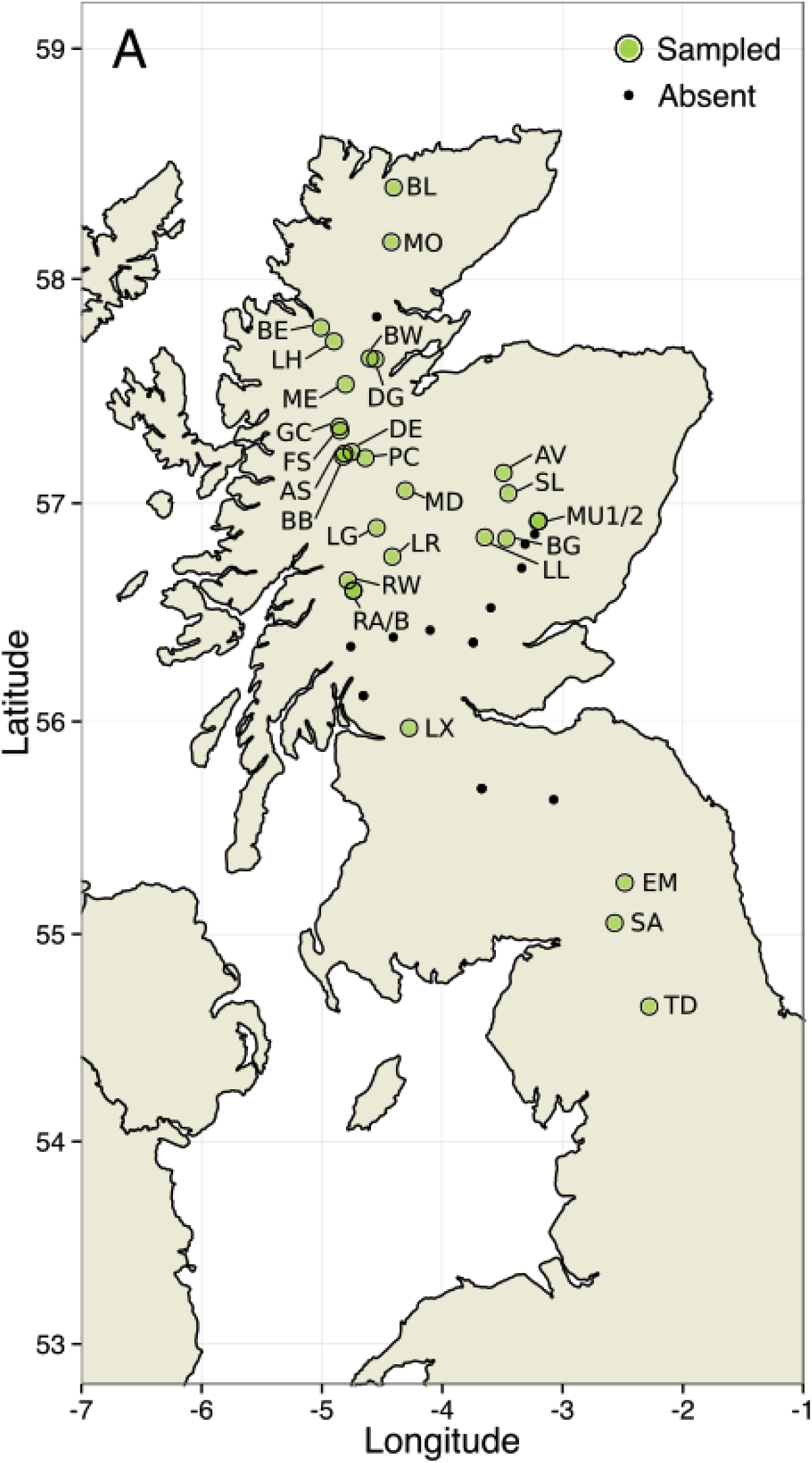
Map of dwarf birch sampling locations in the UK. Adapted from (Borrell et al., 2018).

**Figure S3.**
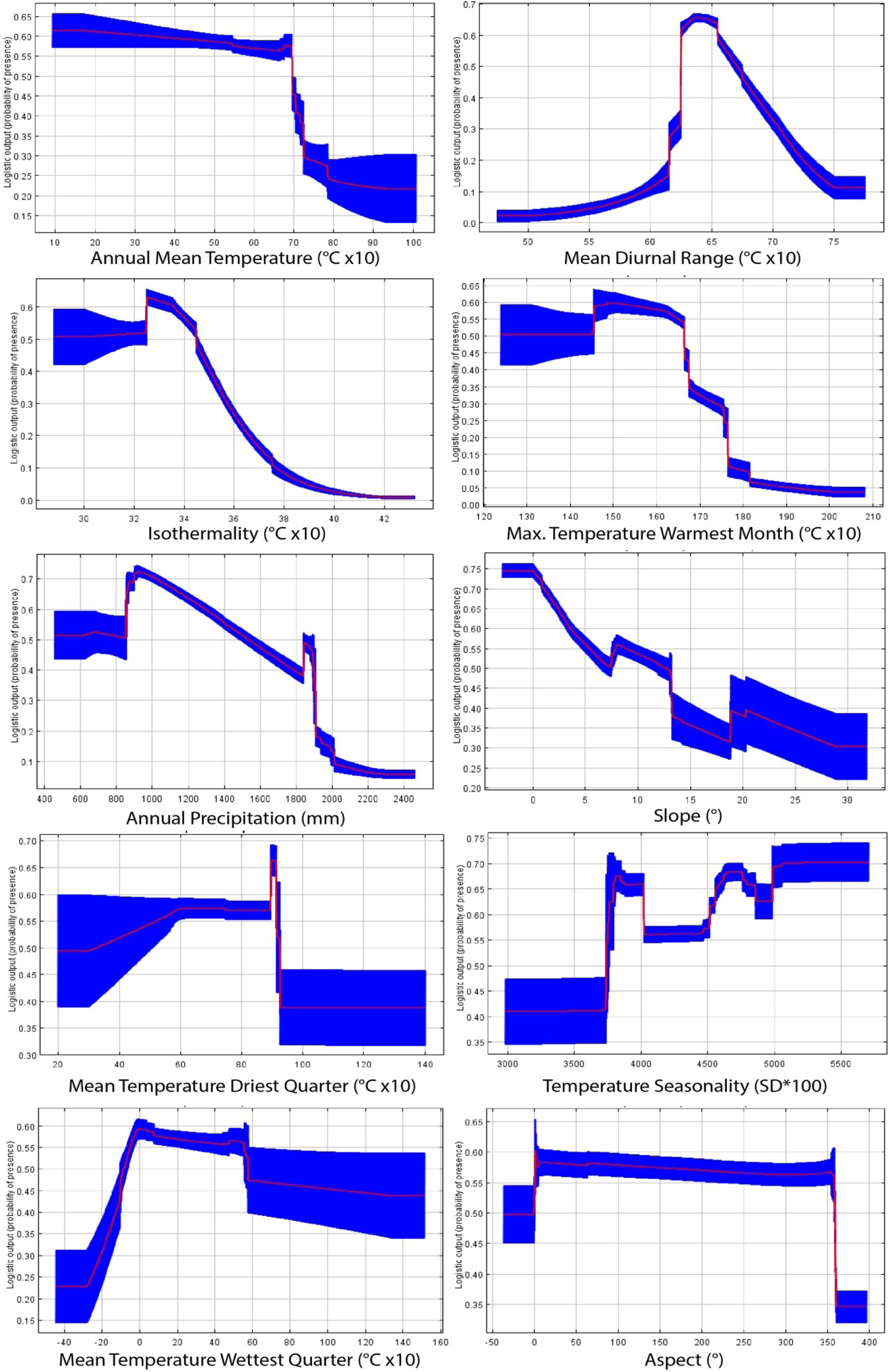
Environmental niche model variable response curves for the 10 retained environmental variables used in this study.

**Figure S4.**
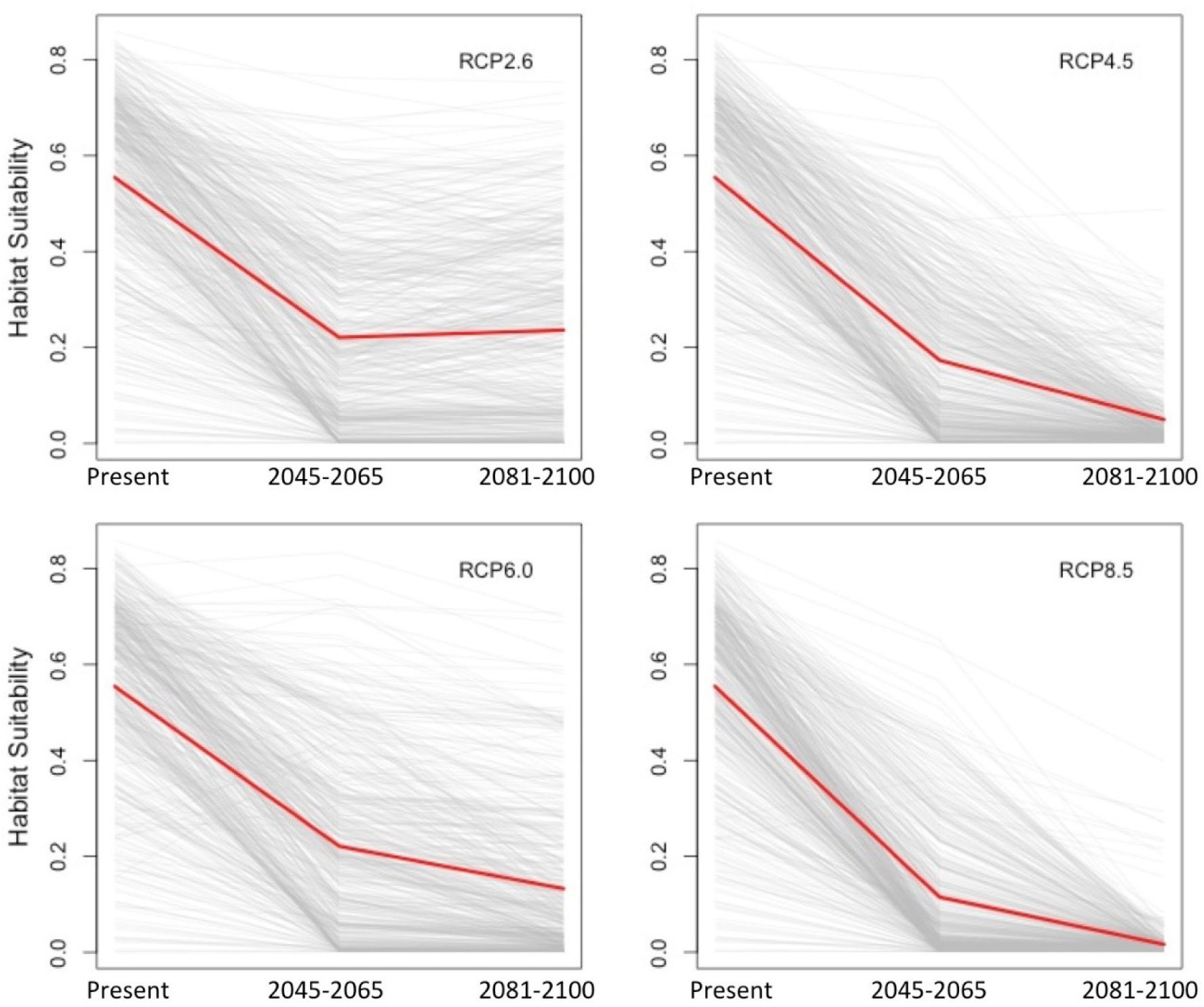
Changes in environmental niche model derived habitat suitability index (HSI) for dwarf birch under four future climate scenarios. Red line indicates overall mean for all recorded locations

**Figure S5.**
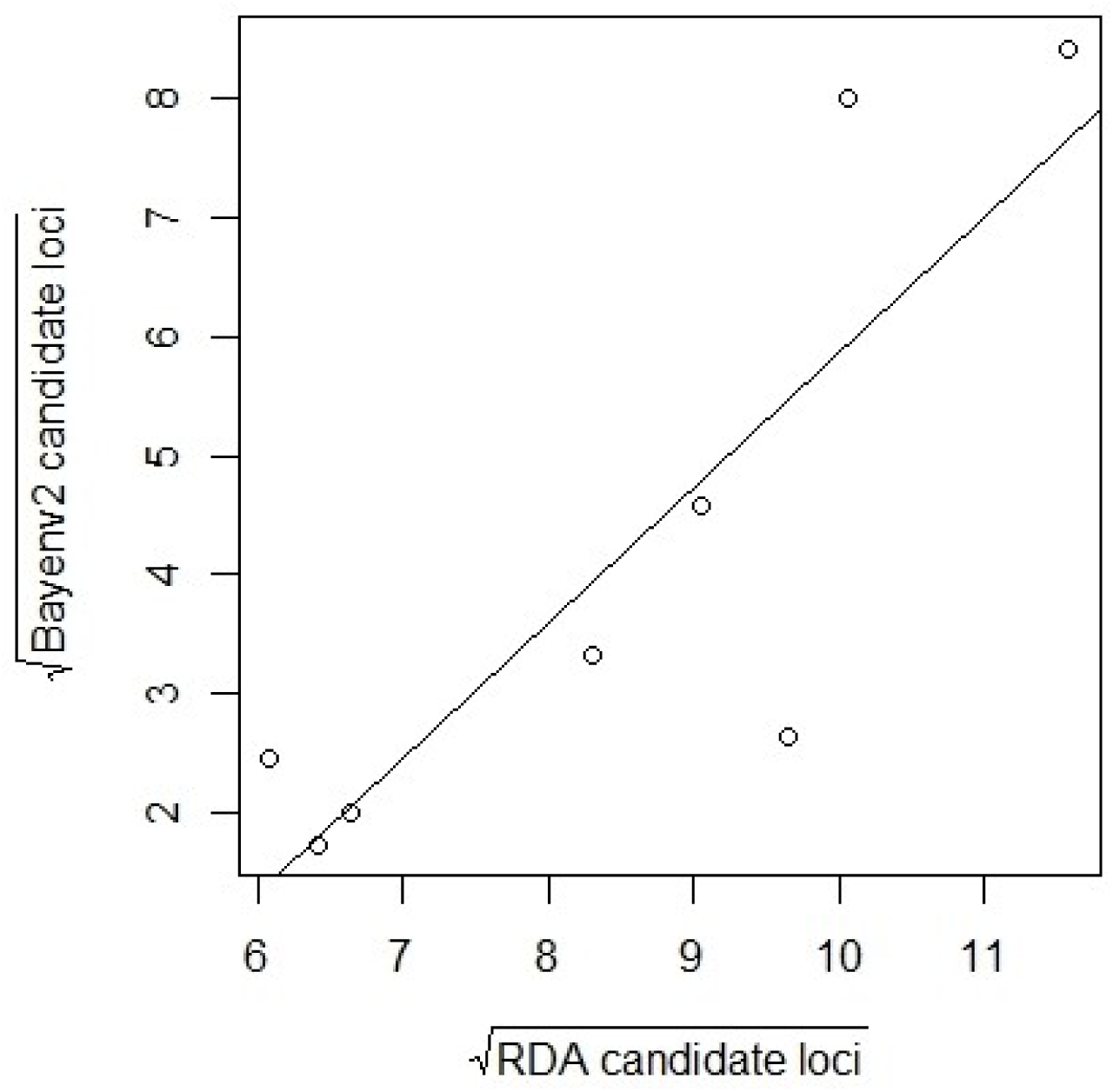
Comparison of the number of the number of candidate adaptive loci identified for each environmental predictor variable in RDA and Bayenv2 GEA analyses.

**Figure S6.**
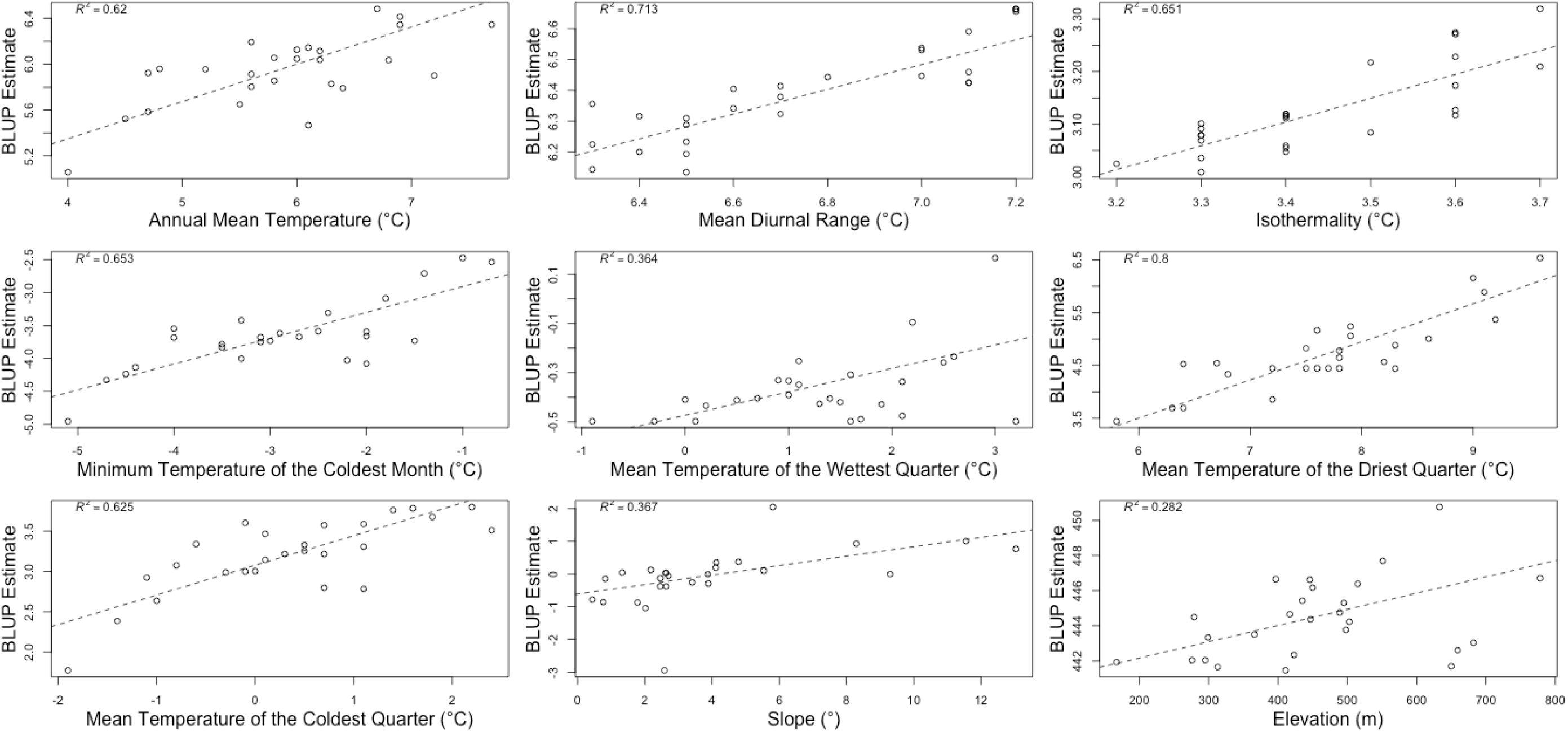
Genotype-environment association plots for nine environmental variables each with more than six associated loci, with dotted line denoting theoretical optimum genotype.

**Figure S7.**
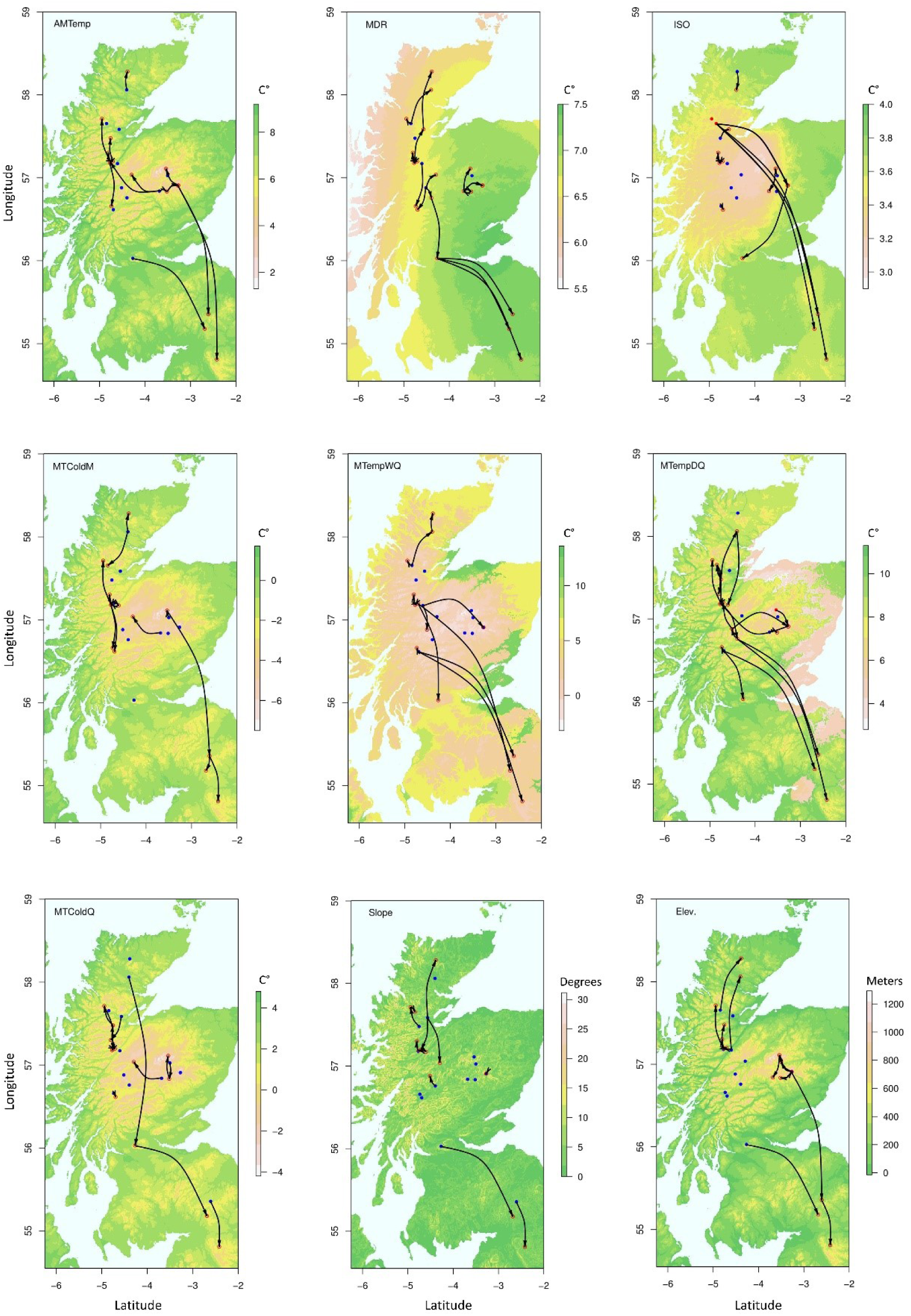
Assisted gene flow maps for nine environmental variables with more than six significantly associated loci.

